# High-resolution lung adenocarcinoma expression subtypes identify tumors with dependencies on *MET, CDK4, CDK6*, and *PD-L1*

**DOI:** 10.1101/2022.02.20.481157

**Authors:** Whijae Roh, Yifat Geffen, Mendy Miller, Shankara Anand, Jaegil Kim, David Heiman, Justin F. Gainor, Peter W. Laird, Andrew D. Cherniack, National Cancer Institute Center for Cancer Genomics Tumor Molecular Pathology (TMP) Analysis Working Group, Gad Getz

## Abstract

Lung adenocarcinoma is one of the most common cancer types with various treatment modalities. However, better biomarkers to predict therapeutic response are still needed to improve precision medicine. We utilized a consensus hierarchical clustering approach on 509 LUAD cases from TCGA to identify five robust LUAD expression subtypes. We then integrated genomic (patient and cell line) and proteomic data to help define biomarkers of response to targeted therapies and immunotherapies. This approach defined subtypes with unique proteogenomic and dependency profiles. S4-associated cell lines exhibited specific vulnerability to CDK6 and CDK6-cyclin D3 complex gene, CCND3. S3 was characterized by dependency on CDK4, immune-related expression patterns, and altered MET signaling; experimental validation showed that S3-associated cell lines responded to MET inhibitors, leading to increased PD-L1 expression. We further identified genomic features in S3 and S4 as biomarkers for enabling clinical diagnosis of these subtypes. Overall, our consensus hierarchical clustering approach identified robust tumor expression subtypes, and our subsequent integrative analysis of genomics, proteomics, and CRISPR screening data revealed subtype-specific biology and vulnerabilities. Our lung adenocarcinoma expression subtypes and their biomarkers could help identify patients likely to respond to CDK4/6, MET, or PD-L1 inhibitors, potentially improving patient outcome.

**Significance:** Through integrative analysis of genomic, proteomic, and drug dependency data, we identified robust lung adenocarcinoma expression subtypes and found subtype-specific biomarkers of response, including CDK4/6, MET, and PD-L1 inhibitors.

## Introduction

Lung cancer is the most prevalent cause of death from cancer worldwide (1). The two major histological classes of lung cancer are: (i) non-small-cell lung cancer (NSCLC) and (ii) small-cell lung cancer (SCLC). Of these, NSCLC is the most common histological type and is further divided into two major subtypes: lung adenocarcinoma (LUAD) and lung squamous cell carcinoma (LSCC, previously termed “LUSC”). Previous studies classified LUAD into molecular subtypes based on genomic (2, 3, 4) and proteogenomic (5) profiling of tumors and then associated these subtypes with clinical outcomes. Two of the largest published studies on LUAD subtypes are: (i) the original The Cancer Genome Atlas (TCGA) LUAD subtyping paper published in 2014 that used 230 patients (the largest number of patients available at the time) to identify three subtypes based on mRNA expression –– Proximal Inflammatory (PI), Proximal Proliferative (PP), and Terminal Respiratory Unit (TRU) (2); and (ii) the TCGA Pan-Lung study in 2017 that analyzed both the LSCC and LUAD cohort and identified 8 subtypes, 6 of which are enriched with LUAD tumors (4). To date, the TCGA LUAD cohort has increased to a total of 509 LUAD cases, offering increased power to identify higher resolution subtypes, further refining the LUAD subtypes defined in the original TCGA study.

Robust LUAD subtyping can substantially aid in determining the most effective therapies that target subtype-specific vulnerabilities. Thus far, molecular therapies for LUAD have focused on targeting various genomic alterations, such as the RAS/RAF/RTK pathway. These include therapies targeting *EGFR, ALK*, and *ROS1* alterations, as well as the more recently approved therapies such as those, targeting *MET, RET, NTRK1*/*2, BRAF* kinases and KRAS^G12C^ mutations (6, 7). Moreover, additional therapies are currently still under development such as ERBB2 inhibitors (8). Recently, immune checkpoint blockades have been approved to treat lung cancer, including inhibitors for PD-1 (pembrolizumab and nivolumab) and PD-L1 (atezolizumab and durvalumab). Previously reported biomarkers of response or resistance to immunotherapy in LUAD include PD-L1 expression (9, 10), tumor mutational burden (TMB) (11, 12, 13, 14), mismatch repair deficiency/microsatellite instability (15), and *STK11* mutation (16). Even with these available therapies, most LUAD tumors continue to progress on therapy, underscoring the need for novel therapeutic approaches.Therefore, more precise and robust subtyping of LUAD tumors and their association with specific treatments can help improve patient prognosis and outcome.

In this study, we integrated multiple data sets: (i) the full 509 LUAD patient cohort in TCGA; (ii) vulnerability data in LUAD cell lines from the Cancer Cell Line Encyclopedia (CCLE) (17, 18) and the Dependency Map (DepMap) (19) repositories; and (iii) proteomic data from the Clinical Proteomic Tumor Analysis Consortium (CPTAC) cohort of LUAD patients (5) to more precisely define therapeutically relevant LUAD subtypes (**Figure 1A**). We show that our analysis indeed yielded distinct subtypes compared with the previously published expression-based subtypes, with higher-resolution partitioning of previously defined subtypes. Moreover, our experimental work *in vitro* links selected subtypes with potential subtype-specific therapeutic targets, and we identify a small number of biomarkers that could be used in the clinic to classify patients into our most clinically relevant subtypes, which could help guide clinical decision making.

**Figure 1.**
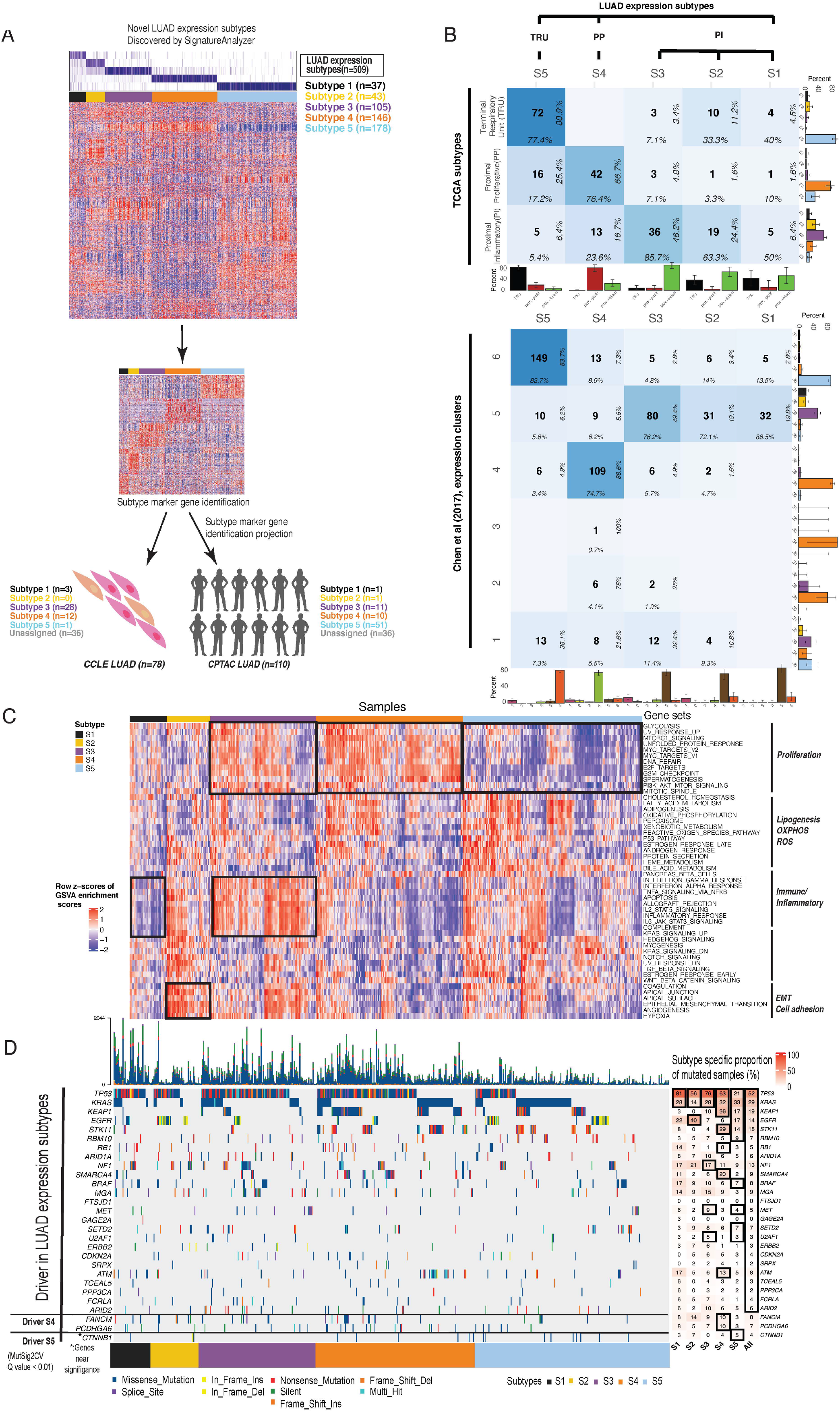
Study design and mutational landscape of LUAD expression subtypes. **(A)** 5 LUAD expression subtypes were identified by *SignatureAnalyzer*. The upper heatmap shows the values of the normalized H matrix identified by the SignatureAnalyzer (row: five expression subtypes, column: 509 TCGA LUAD samples). Samples with normalized association scores higher than 0.6 to a certain subtype were assigned to the subtype. The patient size for each subtype ranged from 7.3% (least common subtype) to 35% (most common subtype) of all cases. The lower heatmap shows the row z-scores of mRNA expression of 100 subtype marker genes for each subtype. Expression subtypes for CCLE LUAD samples (n=78) and CPTAC LUAD samples (n=110) were determined by projecting LUAD expression subtypes to each tumor sample using subtype marker gene expression. Tumor samples were assigned to certain expression subtypes based on normalized association values (cutoff of 0.6). **(B)** The confusion matrix shows concordance between our LUAD expression subtypes and TCGA LUAD expression subtypes (or the Chen et al 2017 expression subtypes (4)). The cell count in the middle shows the number of samples overlapping between two subtypes. The column-wise proportion is shown at the bottom of each cell, and the row-wise proportion is shown on the right side of each cell. The bar plots at the bottom show the column-wise proportion, and the bar plots on the right side of the heatmap show the row-wise proportion. **(C)** The heatmap shows overall pathway activation profiles (in row z-scores of GSVA enrichment scores for MSigDB hallmark gene sets) in LUAD expression subtypes. **(D)** Driver mutations identified by MutSig2CV (point mutations, indels; *Q* value < 0.01) for each LUAD expression subtype and their associated proportion.

## Methods

### Data availability

Publicly available data generated by others were used by the authors. TCGA LUAD expression matrix was obtained from the PanCanAtlas study (20). Survival data for TCGA LUAD samples was obtained from the integrated TCGA pan-cancer clinical data (28). The omics data and CRISPR knockout data for CCLE LUAD cell line samples were obtained from the Dependency Map (DepMap) portal (https://depmap.org/portal/; DepMap Public 21Q2 dataset) (19). Data used in this publication were generated by the National Cancer Institute Clinical Proteomic Tumor Analysis Consortium (CPTAC). Genomics and proteomics data for CPTAC LUAD samples were obtained from the previous study (5). Proteomics datasets were obtained from the CPTAC data portal for LUAD:
https://cptac-data-portal.georgetown.edu/study-summary/S054. Proteomics data processing was performed as previously described (5).

### Ethics approval and consent to participate

All datasets analyzed are publicly available. Ethics approval and consent were obtained in the original papers as required

### TCGA LUAD expression matrix

Batch-corrected upper quartile normalized RSEM (RNA-Seq by Expectation-Maximization) data for TCGA LUAD cohort from the PanCanAtlas study (20) was used for analysis.

### Identification of TCGA LUAD expression subtypes and subtype labeling in CCLE and CPTAC LUAD samples

For expression subtyping, BayesianNMF (21) with a consensus hierarchical clustering approach was applied to the log_2_(RSEM) TCGA LUAD gene expression data as described previously (22), (23, 24). Expression subtype classifiers were then derived as previously described (23). The association of samples from either CCLE or CPTAC RNA-seq samples to the TCGA LUAD expression subtypes (normalized **h**_new_ matrix) was determined by modeling the gene expression matrix of CCLE/CPTAC RNA-seq samples **X**_new_ conditioned on **W**\*_TCGA_ to best approximate **X**_new_ ∼ **W**\*_TCGA_ **h**_new_ for the differentially over-expressed subtype markers (100 marker genes in each subtype) in TCGA LUAD expression subtypes. CCLE and CPTAC RNA-seq samples were assigned to one of the five identified TCGA LUAD expression subtypes if the normalized association (normalized **h**_new_ matrix) with one of the TCGA subtypes was larger than 0.6 (the cutoff of 0.6 instead of 0.5 was used to be more conservative).

### Mutation significance analysis

MutSig2CV (25, 26) was applied to identify significantly mutated genes, and GISTIC 2.0 (27) was applied to identify significant focal copy number alterations in a cohort of samples of interest (all TCGA LUAD samples, each of five TCGA LUAD expression subtypes). Gene amplification in TCGA LUAD was based on the entries having values of +2 (high-level threshold) or +1 (low-level threshold) in the ‘all_thresholded.by_genes.txt’ from GISTIC 2.0. Gene deletion in TCGA LUAD was based on the entries having values of -2 (high-level threshold) or -1 (low-level threshold) in the ‘all_thresholded.by_genes.txt’ from GISTIC 2.0. Gene amplification and deletion in CCLE LUAD was based on a log2 copy number ratio threshold of 0.3. Due to small sample size of CPTAC LUAD cohort (n=1 for S1, n=2 for S2, n=13 for S3, and n=13 for S4), MutSig2CV and GISTIC 2.0 could not be applied for CPTAC LUAD cohort. As an alternative, the proportion of samples with recurrent SCNAs in the TCGA LUAD cohort with those in the CPTAC LUAD cohort was compared.

### Pathway analysis

Single-sample gene set variance analysis (GSVA) was performed on the log_2_(RSEM) TCGA LUAD gene expression data and CPTAC LUAD gene expression data (TPM) using the gsva function (method=“gsva”, mx.diff=TRUE) from the R package ‘GSVA’ (v.1.30.0). GSVA implements a non-parametric method of gene set enrichment to generate an enrichment score for each gene set within a sample. The Molecular Signatures Database (MSigDB) gene sets v.6.1 were used to represent broad biological processes. The pathways with significantly different activities across the subtypes were identified based on (i) FDR-adjusted *P* value < 0.05 and (ii) mean difference of GSVA enrichment scores between subtypes of interest vs. others > 0.2 or < - 0.2.

### Survival analysis

Disease-specific survival information of TCGA LUAD patients (‘DSS’: disease-specific survival event, ‘DSS.time’: disease-specific survival time) and other clinicopathologic variables were obtained from an integrated TCGA pan-cancer clinical data resource (28). Kaplan-Meier curves (with the log-rank test *P* values) were plotted using the Surv function in the R package ‘survival’ (v.2.43-1).

### Biomarker analysis

Biomarker discovery was performed by applying lasso logistic regression on either gene expression data or RPPA data (level 4 RPPA data were obtained from the Cancer Proteome Atlas Portal) from the TCGA LUAD cohort (randomly split into 80% training data and 20% test data) to predict subtypes of interest (S3 vs. others or S4 vs. others). For gene expression data, 100 subtype marker genes were used (**Table S5**) as the potential features to test. The best lambda value was chosen to minimize the prediction error rate using the cv.glmnet() function in the R package ‘glmnet’ (v.4.1-1). Threshold values from 0.1 to 1 in increments of 0.1 were tested for the best threshold selection that maximizes AUC values. Accuracy of the model was based on the agreement of the predicted subtypes and the true subtype label in the test data. To reduce the number of features down to five for the 5-feature models, we forced the model to reduce the number of features down to five by increasing the lambda value that controls the amount of the coefficient shrinkage.

### Antibodies and reagents

The following antibody was used for immunofluorescence staining: Recombinant Alexa Fluor® 488 Anti-PD-L1 antibody (ab209959). DAPI was used for nuclear staining (10236276001; Sigma-Aldrich), C-Met inhibitor tivantinib was purchased from Selleck Chemicals (Houston, TX, USA), CDK4/6 inhibitor Palbociclib (PD 0332991 isethionate) was purchased from Sigma-Aldrich. CDK4/6 Inhibitor IV (CAS 359886-84-3) was purchased from Calbiochem.

### Cell cultures

LUAD cell lines (HCC78, HCC827, NCIH1975, NCIH1838,NCIH1395, NCIH1833, NCIH1755,ABC1, CALU3) were obtained from the Meyerson lab. Tests for mycoplasma contamination were negative. Cells were maintained in RPMI-1640 medium supplemented with 10% fetal bovine serum and 1% penicillin-streptomycin.

### Proliferation assay

Cells were seeded in duplicate (1 × 10^4^ in 96 well plates) and treated with DMSO, tivantinib (3µM), CDK4/6 inhibitor (Palbociclib-CDK4 concentration - 11nM; CDK4/6 concentration - 16 nM), or CDK4/6 Inhibitor IV (CINK4; CDK4-specific concentration -1.5 µM). The media and drugs were replenished every 2-3 days. Continuous cell growth was monitored in 96-well plates every 3 hrs for 4 days using the IncuCyte Kinetic Imaging System. The relative confluency was analyzed using IncuCyte software. The reported response percentage for each cell line was calculated as the percent of confluency compared to a DMSO-treated control counterpart. Proliferation assays were repeated 4 times.

### Immunofluorescence microscopy

Cells were seeded in duplicate (5 × 10^4^ in 24 well plates), treated with DMSO or tivantinib, and grown for 2-3 days. Cells were then fixed in 4% paraformaldehyde for 10 min and washed twice in cold PBS. Fluor® 488 Anti-PD-L1 antibody was added for 1 hr incubation in a light-protected environment at room temperature followed by staining the nuclei with DAPI. Fluorescence images were captured using Invitrogen™ EVOS™ FL Imaging System by Thermo Fisher Scientific. The increase in fluorescence was further quantified using ImageJ software.

### Statistical analysis

Statistical analysis was performed using R. Statistical tests included a two-sided Wilcoxon rank-sum test and Chi-squared test.

## Results

### Genomic characterization reveals five LUAD expression subtypes

Since previous studies showed that expression data are the most predictive genomic features of cancer dependencies (19), we first applied a consensus clustering approach, using Bayesian Non-Negative Matrix Factorization (BayesianNMF) to the expression data representing the 509 LUAD cases from TCGA, similar to how we previously identified molecularly distinct subsets of bladder cancer associated with clinical outcomes (22). Our analysis revealed a robust and detailed structure, yielding five LUAD expression subtypes, designated as S1 to S5 (**Figure 1A, Table S1**).

To further explore our expression subtypes, we compared them to the previously defined LUAD expression subtypes –– PI, PP, and TRU (2). Among the 230 TCGA LUAD tumors with information available for these three subtypes, we found that S5 was most closely related to the TRU subtype (77.4% of S5 tumors were from the TRU subtype, and 80.9% of TRU subtype mapped to S5 [Fisher’s exact test *P* = 3.3×10^−24^]), and S4 was enriched with the PP subtype (76.4% of S4 were PP; *P* = 9.4×10^−9^). The S1, S2, and S3 subtypes mostly matched the PI subtype, suggesting that the PI subtype can be further split into three subgroups. Of these, S3 was the most enriched with the PI subtype (85.7% of S3 tumors matched to PI; *P* = 1.7×10^−14^) (**Figure 1B**). We also compared our expression subtypes to the 6 LUAD-enriched mRNA expression clusters from the more recent TCGA Pan-Lung study (4) We found a high concordance across the two TCGA studies and our analysis (**Figure 1B, Table S2**). Note that the Pan-Lung study also defined subtypes using cluster-of-clusters analysis (COCA) that are based on different genomic features, and the level of concordance with these subtypes was lower than with the mRNA-based subtypes ((4); **Figure S1A, Table S2)**.

To explore the biological differences among our five subtypes, we calculated pathway activity levels for each LUAD sample using single-sample gene set variance analysis (GSVA) on the Molecular Signatures Database (MSigDB) hallmark gene sets in order to identify the pathways with significantly different activities across the subtypes. S1 showed a low immune/inflammatory signature, and S2 showed high activity of gene sets associated with epithelial–mesenchymal transition (EMT) and cell-adhesion. Both S3 and S4 showed increased proliferation signatures (*Q* values < 2.5×10^−10^), but only S3 showed high immune/inflammatory signatures (*Q* values < 3.7×10^−17^). S5 distinctively showed low proliferation signatures (**Figure 1C, Table S3**).

To further support the partitioning of the 60 tumors that originally were assigned to the PI subtype and were further partitioned into our S1-S3 subgroups, we searched for differences in pathway activities among them. We found a consistent set of differentially active pathways as determined when using all the samples (e.g., low immune/inflammatory signature in S1; high EMT in S2; high E2F, MYC targets, G2M markers, and interferon alpha/gamma response in S3; Methods), demonstrating that the differences between the S1-S3 subgroups already existed in the original TCGA cohort but likely not detected due to the small sample size (**Figure S1D, Figure 1C, Table S3**). Altogether, we revealed novel, biologically distinct subtypes that were previously grouped together within the single PI subtype.

To associate each of our five expression subtypes with driver events (point mutations, indels, and copy-number alterations), we applied *MutSig2CV* (25) and *GISTIC 2*.*0 (27)* to the tumors in each of the subtypes (**Figure 1D, Figure S2A, Figure S2B**). Consistent with the original TCGA paper (2), we found that S5 (mapping to the TRU subtype) was enriched with *EGFR* mutations/amplifications and S4 (mapping to PP) was enriched with activating *KRAS* and inactivating *STK11* mutations (2). Interestingly, we found that S2 (one of the subgroups of the PI subtype), was also enriched with *EGFR* mutations/amplifications, even with a higher frequency than S5 (40% vs 17%, *P* = 0.0023). S3, a different PI-subgroup, exhibited amplification of *CD274* (PD-L1) (near significance: *Q* value = 0.102), *MET*, and *CDK4*.

Beyond the previous associations of driver events with the S4/PP and S5/TRU subtypes, our analysis of a larger dataset enabled identification of significantly recurrent *SMARCA4, ATM, FANCM*, and *PCDHGA6* mutations as well as amplification of *MET, FGFR1*, and *PIK3CA* in S4; and *BRAF, SETD2*, and *CTNNB1* recurrent mutations in S5 (**Figure 1D, Figure S2A, Figure S2B**). *STK11* mutations in particular were enriched in *KRAS*-mutant S4 tumors (21 *STK11*-mutant tumors among 46 *KRAS*-mutant S4 tumors) (Fisher’s exact test *P* = 0.0029), suggesting that S4 tumors might be more resistant to PD-1 inhibitors (16, 29).

Next, we leveraged 16 previously calculated genomic features for TCGA samples (30) to further explore differences across the S1-S5 subtypes (**Figure S3**). We observed that S2 and S5 (both significantly enriched with EGFR mutations) have lower overall somatic tumor mutation burden (TMB) than the other subtypes, including significantly lower frequencies of both nonsilent mutations and indels, which influence the number of predicted neoantigens (**Figure S3A-D**). In features related to somatic copy-number alterations (SCNAs), we observed that S1 and S4 have significantly higher number of copy-number segments and fraction of genome altered by SCNAs, as well as higher levels of homologous recombination defects and aneuploidy score (**Figure S3E-H)**. These genomic differences provide orthogonal evidence for splitting the prior PI subtype into S1 (higher SCNAs) and S2 (lower TMB) subtypes, with the remaining PI-like tumors falling into S3. Finally, we associated our subtypes with immune cell populations that were derived by Thorsson et al. (30) using CIBERSORT (31) for deconvolving expression data (**Figure S4A**). We found that S2 showed significantly higher TGF-beta levels and a higher fraction of M2 macrophages (**Figure S4B**), which may be explained by secretion of TGF-beta by M2 macrophages to promote immune suppression in S2 (32). Collectively, the genomic characterization of our five expression subtypes shows that each subtype has distinct biology.

Next, we tested whether our expression subtypes were associated with outcome and found a marginally significant association between our subtypes and disease-specific survival (DSS) (*P* value = 0.046). Our S5, which is enriched with the TRU subtype, showed the best prognosis (**Figure S1E**). This result is consistent with the previous TCGA study showing that the TRU subtype had the best prognosis among their three expression subtypes (2).

### Subtype-specific cancer vulnerabilities

We next asked whether the CCLE (17, 18) and DepMap (19) resources, which provide expression data as well as CRISPR and drug screening data for ∼1,100 cell lines, could be leveraged to find subtype-specific cancer vulnerabilities. We first probabilistically classified the 78 CCLE LUAD cell lines with expression data (**Table S4**) into the LUAD expression subtypes (45 CCLE LUAD cell lines have both expression and CRISPR data) using subtype-specific marker genes (**Figure 1A, Figure 2A, Figure S1F, Table S5**, Methods). Since only S3 and S4 were assigned a sufficient number of cell lines (31 and 16, respectively), we focused our downstream analysis on these subtypes. To validate our subtype classification, we confirmed that the S3- and S4-associated cell lines harbored genetic events, somatic point mutations, and copy-number alterations (**Figures S5A and S5B**) that were consistent with patients associated with S3 and S4. We then compared the cancer vulnerabilities of 21 LUAD driver oncogenes between S3/S4 and the other CCLE LUAD cell lines (**Figure 2B**). The analysis did not yield significant S3 vulnerabilities (**Table S6**), but did identify two significant vulnerabilities for S4: *CDK6* and the CDK6-cyclin D3 complex gene, *CCND3* (significant both within the 45 LUAD cell lines and all 114 lung cancer cell lines with both CRISPR and expression data available) (**Table S6**). This finding suggests that S4 tumors may be dependent on the CDK6 pathway and thus potentially vulnerable to CDK6 inhibition.

**Figure 2.**
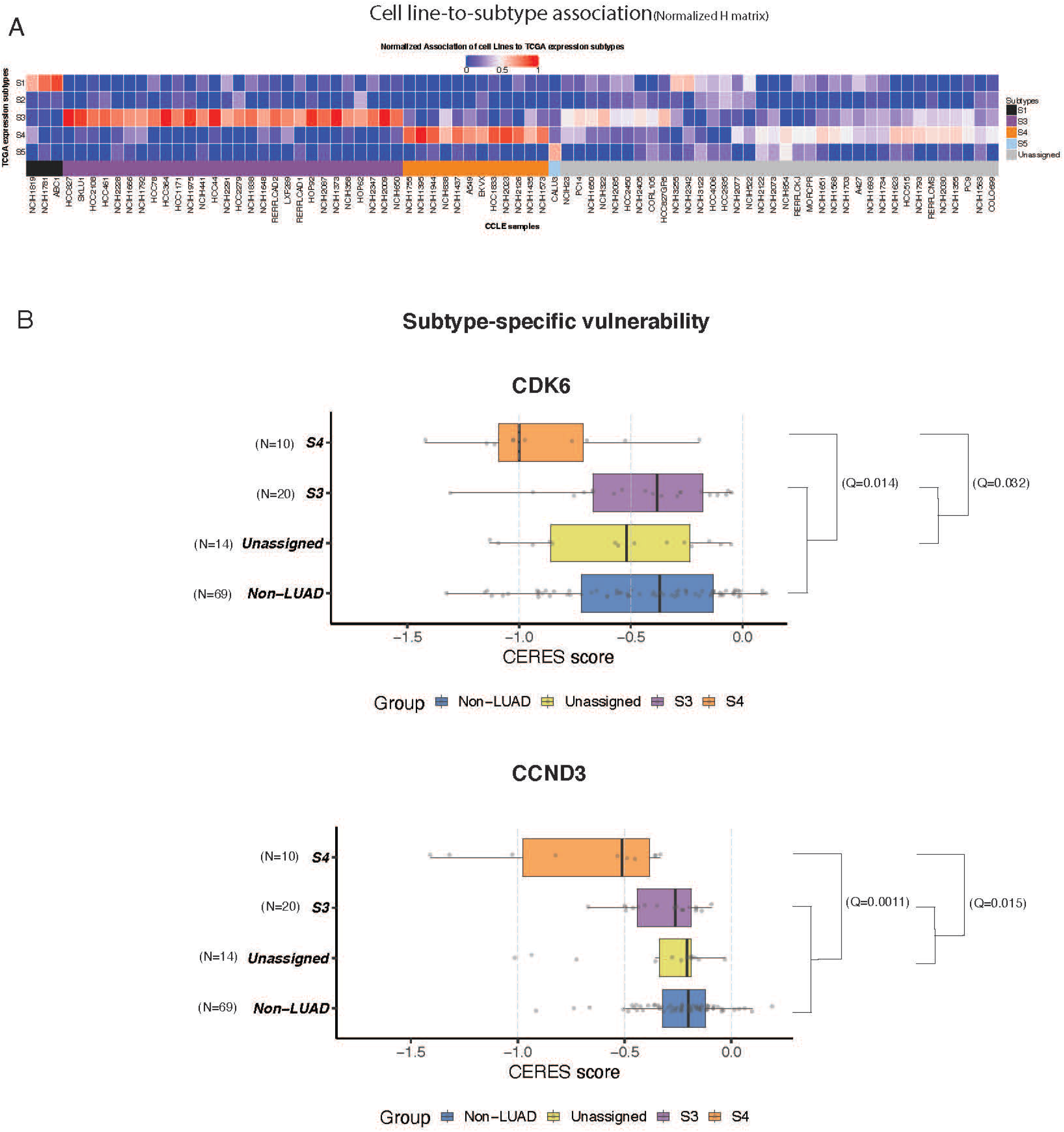
Subtype-specific cancer vulnerabilities. **(A)** The heatmap shows the values of the normalized H matrix. Samples were assigned to the subtype based on two criteria. First, samples should have normalized association scores higher than 0.5 to the subtype. Second, the difference between the highest association score (higher than 0.5) and the second highest association score should be larger than 0.2. Subtype 3 (n=31) and 4 (n=16) cell lines were represented in the CCLE LUAD dataset. **(B)** The boxplots show CERES scores of *CDK6* (top panel) and *CCND3* (lower panel) in S4 versus other cell lines. LUAD driver oncogenes (genes with recurrent point mutations, indels, and SCNAs identified from this study) identified from this study (n=21) were tested. Top genes with subtype-specific cancer vulnerabilities were selected as the genes that meet the following two criteria. First, the genes should have a median CERES score lower than -0.5 in S4. Second, the genes should have a median difference in CERES scores of less than -0.2 when comparing between S4 and other cell lines. The common essential genes (Achilles common essential genes; **Table S6**) were filtered out from the top gene list. *P* values were calculated by the Wilcoxon rank sum test. The one sample assigned to S1 is not shown due to the small sample size of S1.

Although we did not find significant CRISPR vulnerabilities associated with S3, we noticed that *CDK4* was nominally significant (*P* = 0.01; *Q* = 0.13) (**Table S6**), which was consistent with the recurrent genomic alterations in *CDK4* (**Figure S2B; Figure S5B**) in S3. We therefore functionally tested the sensitivity of S3-associated cell lines to CDK4 specific inhibition using two CDK4 inhibitors: Palbociclib and CDK4/6 Inhibitor IV (CAS359886-84-3, a triaminopyrimidine compound that acts as a reversible and ATP-competitive inhibitor; abbreviated here as CINK4). Both compounds are known CDK4/6 inhibitors at high concentrations; however, at low concentrations, they are potent CDK4-only inhibitors that induce G1 cell cycle arrest and senescence in retinoblastoma protein (Rb)-proficient cell lines (33). We treated 9 cell lines with either palbociclib or CINK4 –– 4 from the S3 subtype (HCC78, HCC827, NCIH1975, NCIH1838), 3 from the S4 subtype (NCIH1395, NCIH1833, NCIH1755) and two that were not confidently assigned to any subtype (ABC1, CALU3) –– and measured proliferation with and without drug. As expected, the S3 cell lines showed significantly lower proliferation (higher response) compared to the S4 and unassigned cell lines (Palbociclib: *P* = 1.6×10^−5^ and *P* = 4.1. ×10^−6^ respectively [**Figure S5C**, left panel] and CINK4: *P* = 3.5×10^−3^ and *P* = 3.3×10^−3^ [**Figure S5C**, middle panel]). These results show that the S3 subtype depends on CDK4, suggesting that therapy that includes a CDK4 inhibitor may benefit patients with S3 tumors. Since palbociclib inhibits both CDK4 and CDK6 at higher concentrations, we could not test CDK6-only inhibition in S4 cell lines, and higher doses of palbociclib inhibited proliferation in all cell lines (**Figure S5C**, right panel). Taken together, the CRISPR data and drug sensitivity experiments demonstrate specific vulnerabilities related to CDK4 in S3 and CDK6/CCND3 in S4 subtypes.

### Proteogenomic analysis reveals distinct protein regulation between S3 and S4

To further characterize the expression subtypes at the proteomics level, we first classified the CPTAC LUAD samples (5) to S1-S5 based on the expression of subtype-specific marker genes (**Figure 1A; Figure 3A; Table S7**). Since S3 (n=11), S4 (n=10), and S5 (n=51) were the major subtypes represented in the CPTAC LUAD cohort, we focused our downstream proteogenomic analysis on these subtypes. Consistent with the TCGA data analysis, both S3 and S4 showed increased proliferation signatures, and S3 also showed an increased immune/inflammatory signature (**Figure 3A**). Comparing our expression subtypes with the CPTAC multi-omics clusters (5), we found a good agreement: S3 was enriched with CPTAC multi-omics cluster C1 tumors (PI enriched; 11 C1 subtype tumors out of 11 S3 tumors), S4 was enriched with C3 tumors (PP enriched; 9 C3 subtype tumors out of 10 S4 tumors) (**Figure S6A**), and S5 was enriched with C4 tumors (TRU enriched; 33 C4 subtype tumors out of 51 S5 tumors). As for the TCGA mRNA subtypes, we found a significant overlap with the multi-omic subtypes in CPTAC (5) and evidence of a more refined partitioning of the PI subtype.

**Figure 3.**
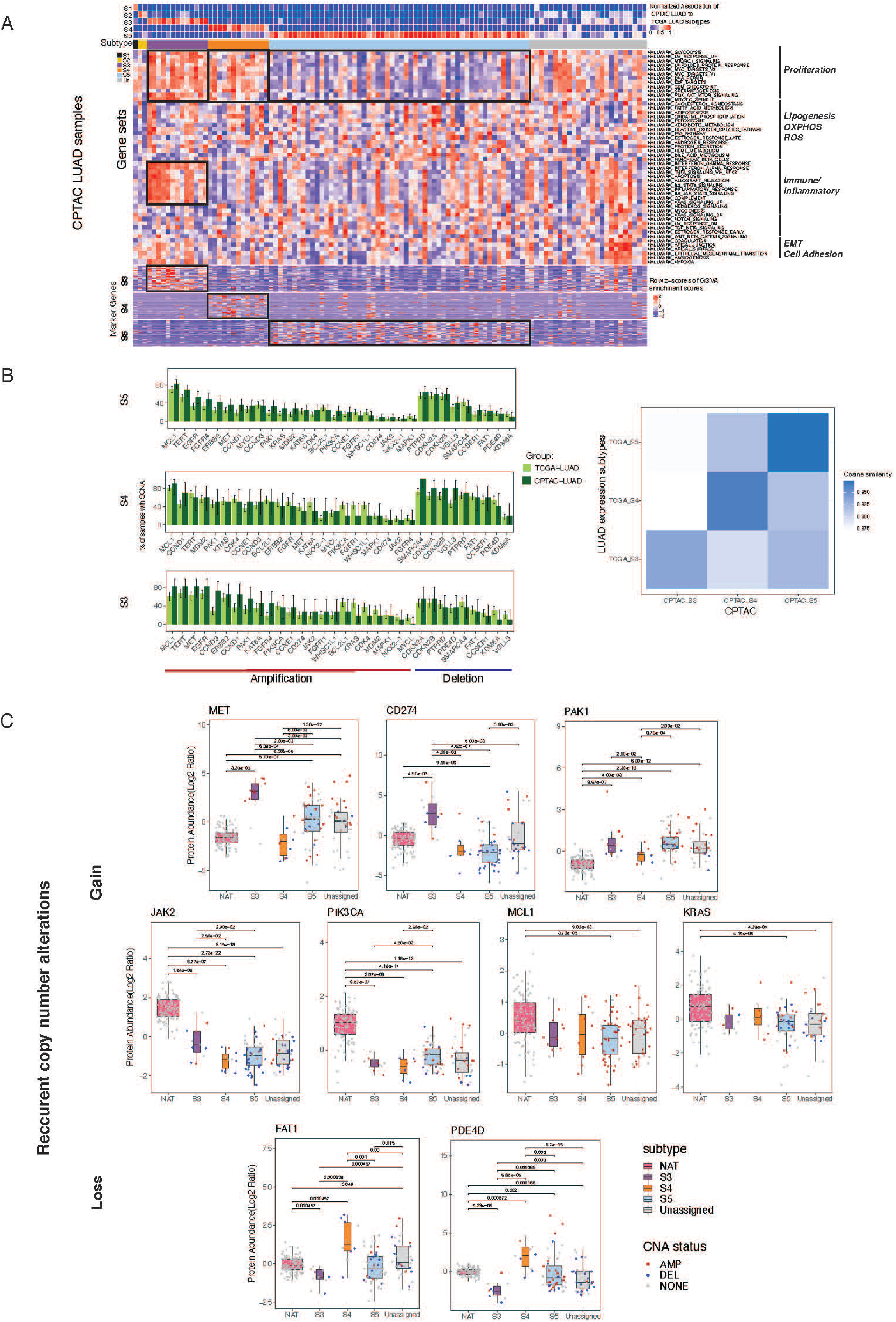
Proteogenomic analysis of genes with subtype-specific recurrent SCNAs. **(A)** The upper heatmap shows the normalized H matrix for CPTAC LUAD samples (row: five LUAD expression subtypes; column: CPTAC LUAD samples). CPTAC LUAD tumor samples were assigned to certain LUAD expression subtypes based on normalized association values (cutoff of 0.6). The column annotation below shows the assigned expression subtypes. The middle heatmap shows the row z-scores of GSVA enrichment scores of MSigDB hallmark gene sets among CPTAC LUAD samples. Pathway activation consistent with LUAD expression subtypes are highlighted with the black box. The lower heatmap shows the row z-scores of S3/S4/S5 marker gene expression across CPTAC LUAD samples. **(B)** The barplots show the proportion of TCGA/CPTAC LUAD samples in S3/S4/S5 with gene amplification (red annotation bar) or deletion (blue annotation bar) for selected genes that had recurrent SCNAs in at least one of the S3/S4/S5 subtypes. The right heatmap shows the cosine similarity among S3/S4/S5 tumors in TCGA and CPTAC data. **(C)** The boxplots show protein abundance of genes with recurrent SCNAs across CPTAC LUAD expression subtypes and their normal adjacent tumors (NAT). Copy number states of genes are shown with different colors (red: amplification; blue: deletion; gray: no SCNAs).

Next, we focused on the genomic and proteomic features of each of our subtypes. The overall frequency profiles of amplification or deletion of significantly copy-number altered genes was similar between the TCGA and CPTAC cohorts for S3, S4, and S5 (cosine similarities > 0.93; **Figure 3B**). Some differences could be attributed to the distinct populations represented in the two cohorts. While the TCGA LUAD cohort is mainly composed of white individuals, the CPTAC LUAD cohort is more diverse, with approximately half white and half Asian individuals (**Figure S6B**). Despite the difference in ethnic background, the fact that the profiles associated with each subtype are more similar to the corresponding subtype than to other subtypes further supports our subtype classification of the CPTAC tumors.

We next explored the effect of recurrent SCNAs on protein expression. Among the genes with recurrent SCNAs (in S3-S5), we compared the available protein expression for these genes across samples classified to S3, S4, S5, and, as controls, also across normal adjacent tissues (NAT) samples and CPTAC LUAD samples that were not confidently assigned (due to low similarity) to any of our subtypes (designated as ‘unassigned’) (**Figure 3C; Figure S6C)**. *JAK2* and *CD274* (PD-L1) showed both recurrent gene amplification and significantly higher protein expression in S3. Interestingly, *MET* showed recurrent gene amplification in both S3 and S4, but its protein expression was significantly up-regulated only in S3 (**Figure 3C**), also exceeding the expression in normal adjacent tissues (*P* < 5.5×10^−4^) (**Figure S6D**). Moreover, S3 tumor samples with *MET* amplification showed much higher *MET* protein expression than S3 tumors with no *MET* amplification, whereas other subtypes showed weaker (or no) correlation between *MET* amplification and *MET* protein expression. Of all the genes that showed recurrent gene deletion in both S3 and S4, only *FAT1* and *PDE4D* also exhibited significant changes to their proteomic expression. Moreover, only S3 exhibited a significantly downregulated protein expression for both FAT1 and PDE4D that was associated with their respective gene loss when compared to both NAT and the other subtypes. We observed a similar trend for mRNA expression in the TCGA LUAD cohort. These findings highlight the need to take into account not only copy-number alterations but also mRNA and protein expression (**Figure S7A**) (34).

### *MET* is a core regulator of proliferation and PD-L1 expression in S3

Our initial pathway activation analysis found that both S3 and S4 upregulate proliferation genes, whereas only S3 highly expresses immune-related genes (**Figure 1C**). To gain additional insight into the underlying biological differences between S3 and S4 (as well as the differences among S1, S2, S5, and the unassigned samples that do not have these two phenotypes), we performed a deeper proteogenomic characterization of our subtypes. First, we noted that *CD274* (PD-L1) showed recurrent gene amplification in S3. We further observed that PD-L1 copy number, mRNA expression, protein expression, and phosphorylation levels were significantly higher in S3 versus S4 (**Figure 3B-C, Figure 4A, Figure S7B**). Since both S3 and S4 showed high proliferation signatures and recurrent *MET* amplification (**Figure 1C, Figure S2B, Figure S7B**), we assessed *MET* copy number and protein expression across subtypes and found that *MET* copy number was significantly higher in S3 versus S4, and that its expression of mRNA, protein, and phosphorylation levels was also higher in S3 versus S4 (**Figure 3B-C, Figure 4B**), echoing the expression pattern of PD-L1. The mRNA and protein expression of *MET* were also significantly higher in S3 versus S4, even when restricting the analysis only to MET-amplified tumors (Q=1.1×10^−8^ for mRNA, Q=2.7×10^−2^) (**Figure S7C**). Additionally, we identified higher MET pathway activation in S3 versus S4 as evidenced by increased phosphorylation levels of GAB1 in S3, a known downstream substrate of MET (**Figure S8A**). Based on a previous study showing a negative correlation between *MET* expression and the expression of T cell effector molecules (granzyme A, granzyme B, and perforin) in the TCGA dataset (35), we asked whether we observe the same pattern in both subtypes. Interestingly, we observed a negative correlation between the expression of *MET* and T cell effector molecules in S3, but not in S4 (**Figure 4C**), suggesting potential immune evasion of S3 tumors associated with *MET* overexpression.

**Figure 4.**
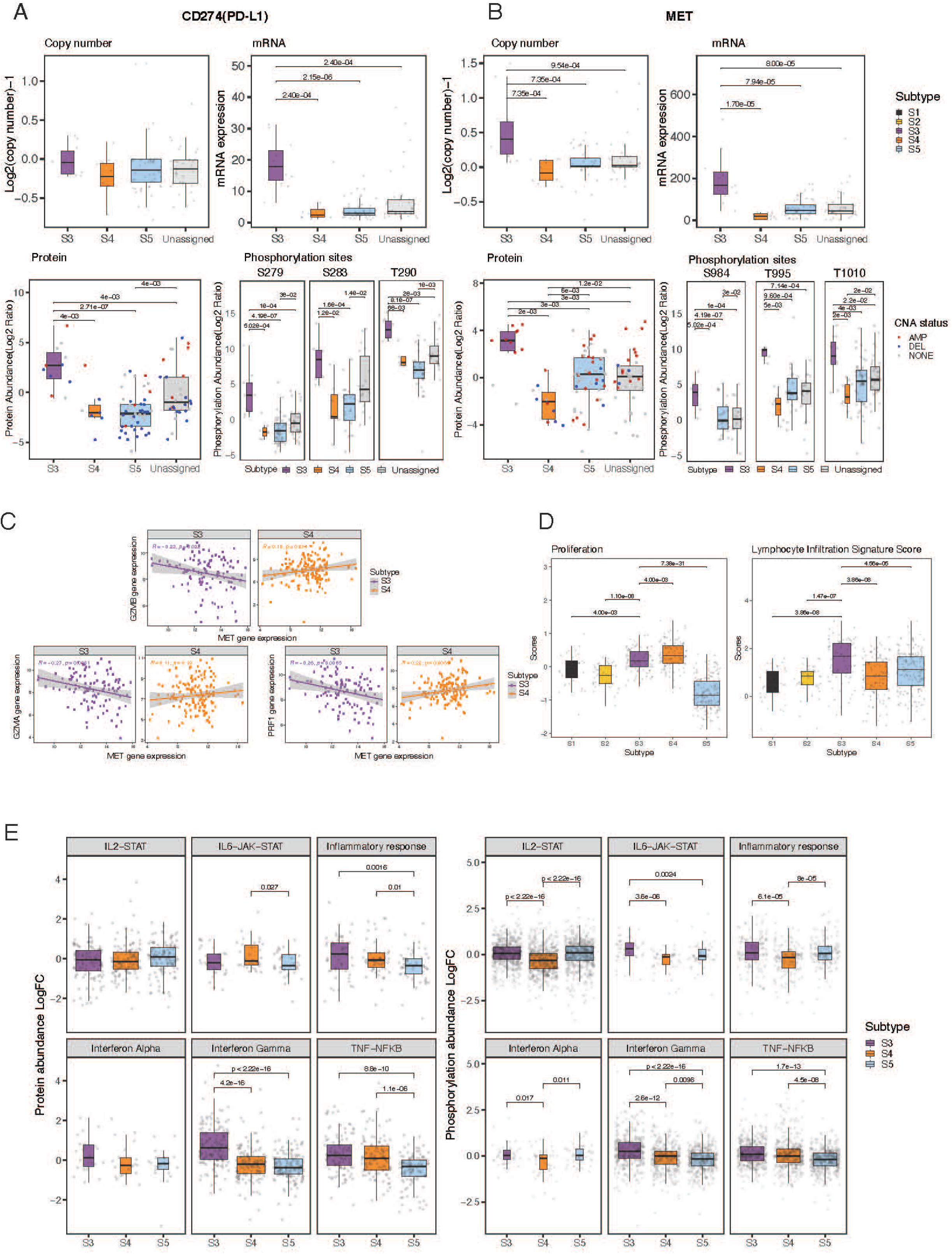
*MET* as a core regulator of proliferation and PD-L1 expression in S3. **(A)** The boxplots show copy number, mRNA expression, protein abundance, and phosphorylation of *CD274* (PD-L1) gene across LUAD expression subtypes in CPTAC data. **(B)** The boxplots show copy number, mRNA expression, protein abundance, and phosphorylation of *MET* gene across LUAD expression subtypes in CPTAC data. **(C)** The scatter plots show correlation between *MET* gene expression and gene expression of cytolytic markers (*GZMB, GZMA*, and *PRF1*) in S3 versus S4. **(D)** The boxplots show proliferation scores and lymphocyte infiltration signature scores (obtained from Thorsson et al., 2018 (30)) across LUAD expression subtypes. **(E)** The boxplots show protein abundance and phosphorylation level of genes in immune-related pathways among CPTAC LUAD S3, S4, and S5.

To next characterize the proteogenomic differences in an alternative way, we evaluated proliferation and immune signatures using previously developed scores for proliferation and lymphocyte-infiltration (30). Again consistent with our results (**Figure 1C**), we found high proliferation scores in both S3 and S4, and a higher immune score only in S3 (**Figure 4D**). Additionally, this alternative characterization method showed that S3 had a significantly higher fraction of anti-tumoral M1 macrophages, suggesting a favorable tumor immune microenvironment for therapy, whereas S4 showed a significantly higher fraction of pro-tumoral Th2 cells (**Figure S4**). We also observed increased interferon-gamma pathway activity in S3 compared to S4 and S5 based on protein expression and phosphorylation data (**Figure 4E**). This finding was further supported by the increased expression of proteins involved in antigen presentation and interferon signaling in S3 (**Figure S8C**). Taken together, these proteogenomic findings support increased immune/inflammatory activity in S3.

To further support and validate our proliferation findings in wet-lab experiments in cell lines, we explored the response of our subtype-specific cell lines (described above) to the MET inhibitor, tivantinib. Tivantinib is a non-ATP-competitive c-Met inhibitor that induces G2/M arrest and apoptosis. We performed a 4-day long proliferation assay to test the response in the different cell lines. S3 showed a significantly increased response (*P* value > 0.001) to tivantinib treatment compared to the other assigned groups (data not shown), and in particular, compared to S4 (**Figure 5A**). Previous reports in NSCLC cells suggested a direct relationship between PD-L1 and MET expression by showing enhanced PD-L1 expression in response to c-MET inhibition (36). To test whether we could also observe this relationship in our subtypes, we assessed PD-L1 levels by performing immunofluorescence staining on the tivantinib-treated cells and controls with an anti–PD-L1 antibody. A significant increase in PD-L1 levels was detected in all subtypes in response to tivantinib (Wilcoxon test *P* value > 0.0001) (**Figure 5B, C**). Since c-MET inhibition has been shown to drive PD-L1 expression by suppressing glycogen synthase kinase 3 beta (GSK3β) (36), we next tested the correlation of mRNA expression between MET and GSK3β in the LUAD cell line data and found a significant positive correlation only in the S3 subtype (Pearson correlation coefficient=0.46, *P* value=0.016) (**Figure 5D**).

**Figure 5.**
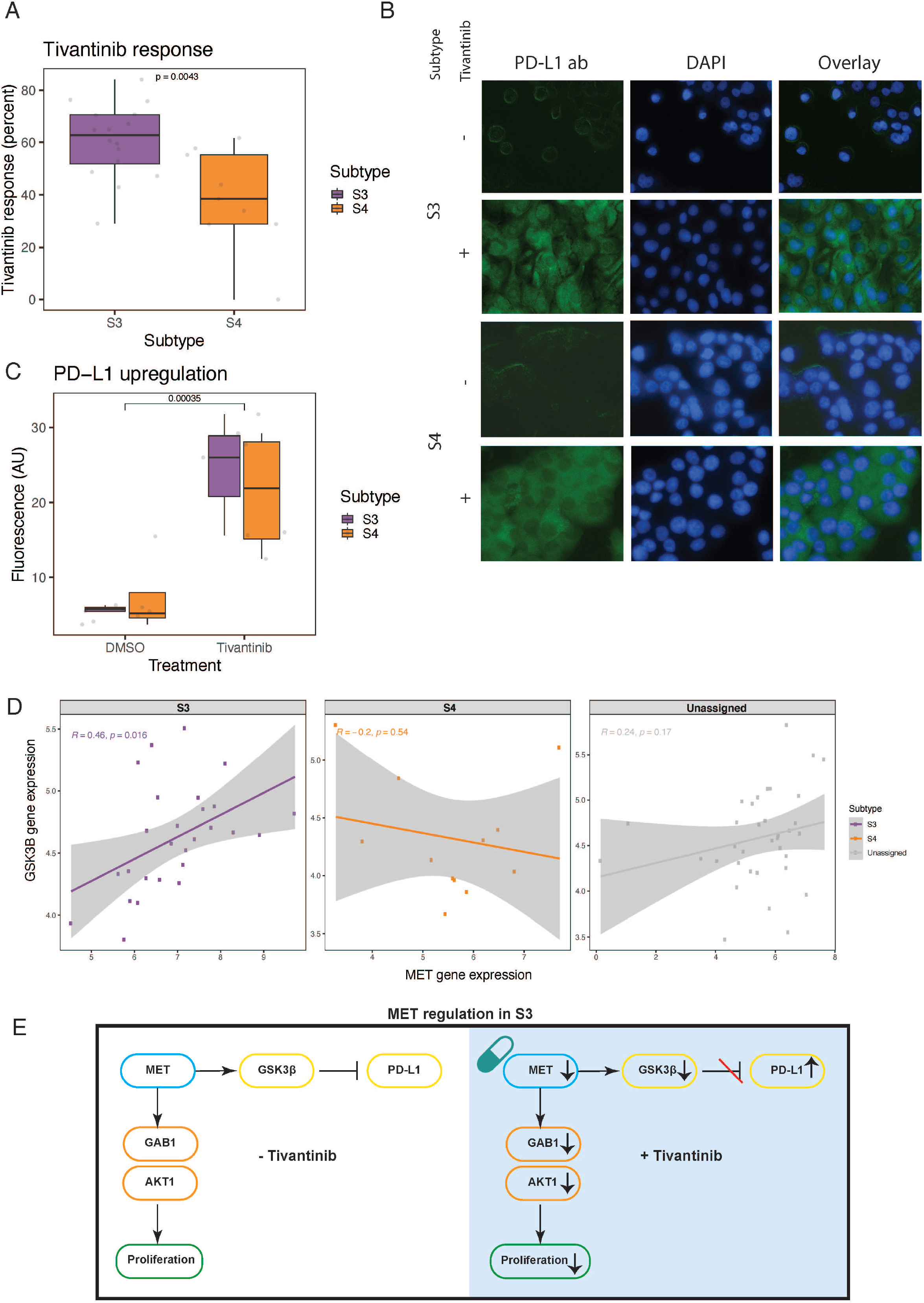
c-MET inhibition drives PD-L1 expression in cell lines. **(A)** The boxplots show response to tivantinib measured by the delta change in confluency between treated and untreated (DMSO only) cell lines within each subtype. **(B)** Immunofluorescence staining under tivantinib treatment. Anti-PD-L1 antibody (Green; first column on the right); DAPI for nuclear staining (Blue; middle column), and overlay of both stainings (left column). **(C)** The boxplots show quantification of fluorescence using ImageJ software after background correction. **(D)** The scatter plots show correlation between *MET* gene expression and gene expression of GSK3β in CCLE data in the different subtypes. **(E)** The schematic diagram shows that *MET* is a core regulator of proliferation and that PD-L1 expression is regulated through the GSK3β axis downstream of *MET* in S3 tumors, both with (right panel) and without (left panel) tivantinib treatment.

Collectively, these data suggest a model for S3 in which MET plays a key role in driving proliferation through GAB1/AKT1, and it can also upregulate PD-L1 expression through the GSK3β axis, potentially for immune escape. Additional synergistic players, found to have higher protein expression in S3 vs S4, such as BCL2L1 and the MCM family, also likely further contribute to the proliferation of cells in S3 (**Figure 5E; Figure S8B**). Hence, S3 tumors may respond to a combined MET and PD-L1 inhibitors.

### Biomarkers for identifying patients with S3 or S4 tumors

Lastly, we were interested in identifying biomarkers for S3 and S4 so that patients with these subtypes could be readily identified in the clinic. For gene-expression data, we used the subtype marker genes defined above (**Table S5**) as the potential features to test. The best prediction model for S3 (23 genes) had an accuracy of 95%, and the best prediction model for S4 (27 genes) had an accuracy of 85% (**Table 1, Table S8**). To identify immunohistochemistry markers, we also considered TCGA reverse-phase protein array (RPPA) data as potential proteomic features. The best prediction models for S3 and S4 contained 20 and 24 protein features, respectively, and were each 91% accurate. Additionally, for better clinical utility, we forced the model to reduce the number of features down to five (Methods). Interestingly, the five-feature model for S3 based on the RPPA data (PD-L1, JAK2, MIG6, P70S6K1, GATA6) still showed a high model accuracy of 91%, whereas the five-feature model for S4 (BIM, CAVEOLIN1, FOXM1, PKCPANBETAII_pS660, NRF2) had a reduced accuracy of 74%. Overall, these results show that we can reach high prediction accuracies for S3 and S4 using both gene expression and RPPA data.

**Table 1.**
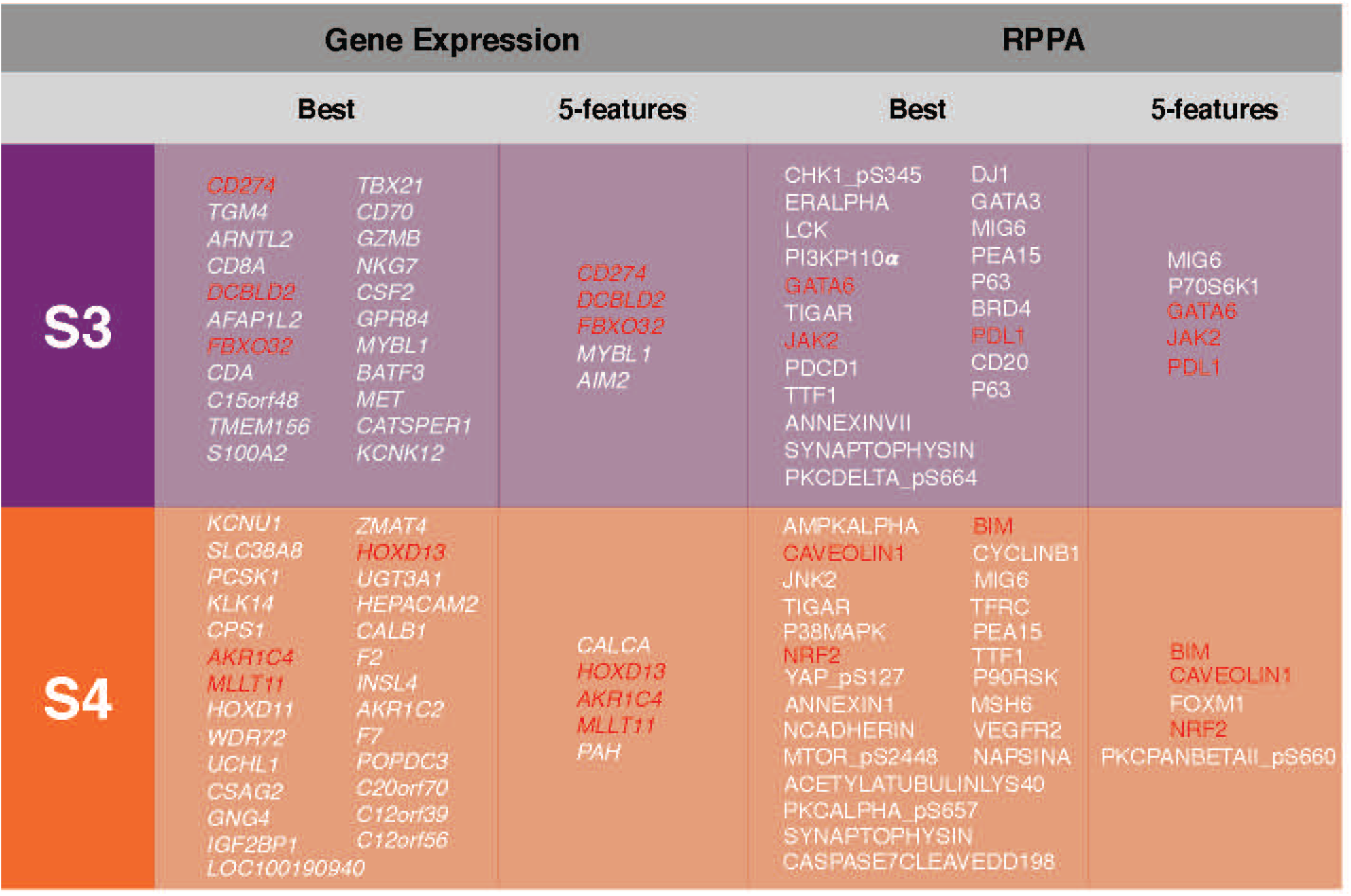
Biomarker discovery for LUAD expression subtypes. Summary table of biomarkers of LUAD expression subtypes S3 and S4 based on gene expression data and RPPA data. The best model represents the model with the lambda that minimizes the cross-validation prediction error rate. The 5-feature model represents the model with the only five features included for parsimony of the model and clinical utility. Features that are included in both the “best models” and “5-feature models” lists are highlighted in red.

## Discussion

Over the past several years, lung cancer subtypes have been studied to reveal new biology associated with clinical outcomes (2, 3, 4, 5). The subtypes identified in these studies were consistent with the PI, PP, and TRU subtypes defined in the original TCGA study, and the subtypes we define here also align well with these original subtypes. Importantly, we were sufficiently powered to further partition the PI subtype into 3 subgroups (S1-S3). By integrating genomic and proteomic data, we further identified distinct biology in each of our subtypes. Notably, the biology of these subtypes was quite distinct from the biology of subtypes identified by the CPTAC LUAD study (5) because the CPTAC LUAD subtypes were based on multi-omics data from a smaller sample size than ours (<5-fold, with only one S1 and no S2 tumors found in the CPTAC LUAD cohort).

We identified multiple subtype-specific significantly recurrent mutations (point mutations, indels, and SCNAs) that the previous TCGA LUAD study (2) was underpowered to detect (**Table 2**). These findings show that the newly identified expression subtypes are associated with distinct tumor biology and might also serve as biomarkers of response to targeted therapies (e.g., response to *EGFR* inhibitors and TGF-beta inhibitors for S2; response to PD-L1, MET, and CDK4 inhibitors for S3; and resistance to PD-1 inhibitors for S4 due to *STK11* mutations (16) (**Table 2**)). Integrative analysis of TCGA and DepMap data also demonstrated the proof-of-concept idea that leveraging the genome-wide CRISPR screening data and expression subtypes of cell lines can identify novel therapeutic targets for specific expression subtypes.

**Table 2.** Subtype-specific proteogenomic features and potential therapeutic targets for subtypes. Summary table of subtype-specific proteogenomic features that were identified from this study, as well as potential therapeutic targets for each subtype. Highlighted features are noted by their respective subtype color - Yellow-S2, Purple-S3, Orange-S4.

|  | Point mutations/<br>Indels | CNAs | Activated<br>pathways | TMB | Signaling | Immune cell<br>subsets | Cancer<br>vulnerability in<br>cell lines | Protein<br>levels | Potential<br>therapeutic targets |
| --- | --- | --- | --- | --- | --- | --- | --- | --- | --- |
| S1 |  |  |  | High |  |  |  |  |  |
| S2 | EGFR mutations | EGFR<br>amplification | EMT,<br>Cell adhesion |  | TGF-beta<br>response | M2 macrophage<br>(pro-tumoral) |  |  | EGFR inhibitors,<br>TGF-beta inhibitors |
| S3 |  | MET, PD-L1<br>amplification | Proliferation,<br>Immune/<br>Inflammatory | High | IFN-gamma<br>response | M1 macrophage<br>(anti-tumoral) | CDK4 | High MET,<br>High PD-L1 | CDK4/MET + PD-L1<br>inhibitors |
| S4 | STK11 mutations | MET, FGFR1,<br>PIK3CA<br>amplification | Proliferation | High |  | Th2 cells | CDK6 |  | CDK6 inhibitors |
| S5 | CTNNB1 mutations |  | Lipogenesis,<br>OXPHOS, ROS |  |  |  |  |  |  |

Our proteogenomic analysis provided additional support for the need to take into account not only copy-number alterations but also mRNA and protein expression when characterizing the biology of the different subtypes (37, 38). For example, our observation that *MET* amplification had a profound impact on its protein expression in S3 but not in the other subtypes suggests that the mRNA and protein expression of these genes may, in some cases, be affected by negative feedback loops or other types of regulation that reduces the effect of the increased DNA copy-number. Collectively, these results highlight the importance of integrating the analysis between genomic and proteomic data to reveal underlying subtype-specific biology (39, 40).

Leveraging proteogenomic data suggest that *MET* amplification in S3 tumors may lead to cell proliferation through the GAB1/AKT1 axis. S3 tumor–specific positive correlation between *MET* gene expression and PD-L1 gene expression also shows that the *MET* gene may regulate PD-L1 expression in S3 tumors through GSK3β, consistent with findings in lung cancer cell lines (35). A recent study demonstrated that MET amplification attenuates immunotherapy response by inhibiting STING in lung cancer and that targeted MET inhibition could increase the efficacy of immunotherapy (41). In our data, the MET–STING axis was attenuated only in S4, but not in S3, suggesting that the MET–GSK3β–PD-L1 axis may play a more important role in S3 than the MET– STING axis. Thus, in S3, MET might be a core regulator of two important cancer-related functions: (i) immune escape by upregulating PD-L1 expression, and (ii) proliferation through a synergistic effect with increased expression of *BCL2L1* and MCM-family members (**Figure S8B**) (42, 43). It is possible that PD-L1 expression–related suppression of anti-tumor immunity may explain the poor performance of the c-MET inhibitor tivantinib in the clinical trial, in addition to it containing mixed tumor subtypes rather than a selected patient population (44). Consequently, combination therapy targeting MET and PD-L1 could be synergistic for S3 tumors. Additionally, since S3 tumors also have relatively high TMB and interferon-gamma gene expression signature, this tumor subtype that accounts for approximately 20% of all LUAD patients (105 out of 509 TCGA LUAD tumors) is likely to respond well to the combination of MET inhibitors and PD-L1 blockade (45).

Since S3 cell lines also had *CDK4* cancer vulnerability and showed high response to *CDK4* inhibitors (**Figure S5C**), S3 tumors might also respond well to combined CDK4 inhibitors and PD-L1 blockade, consistent with the findings from multiple mouse model studies on combined CDK4/6 inhibitors and immune checkpoint blockades (46, 47). Therefore, we suggest that dedicated preclinical studies should be performed in tumor models representing the different tumor subtypes. These findings raise a potential clinical therapeutic hypothesis that membership in the S3 subtype can serve as a biomarker of response to combination immunotherapy targeting *CDK4* or MET together with PD-L1 inhibitors. Since S4 tumors showed recurrent *CCND3* amplification, and S4-associated cell lines showed a significantly stronger dependency on CDK6 (**Figure 2B**), future studies specifically targeting CDK6 in S4 cell lines and patients would also be of interest.

Overall, our study demonstrates that a BayesianNMF approach can identify novel tumor expression subtypes, and that integrative analysis of multi-modal data (genomics, proteomics, and CRISPR screening data) can identify subtype-specific biology and vulnerabilities. Generation of mouse models representative of our LUAD expression subtypes would allow *in vivo* experimental validation of drug response associated with each subtype (46, 47, 48). Since expression subtypes can represent both the tumor cells and their microenvironment –– both of which can contribute to treatment response or resistance –– they can potentially identify more clinically relevant tumor subtypes. Future studies, at the single-cell level, could decouple the contribution of different cell types and potentially reveal new subtype-specific biology as well as cell types and states associated with clinical outcomes (47, 49, 50).

### Consortia

The participants in the National Cancer Institute (NCI) Center for Cancer Genomics (CCG) Tumor Molecular Pathology (TMP) Analysis Working Group are: Jean C. Zenklusen, Anab Kemal, Ina Felau, John A. Demchok, Liming Yang, Martin L. Ferguson, Roy Tarnuzzer, Samantha J. Caesar-Johnson, Zhining Today Wang, Rehan Akbani, Andre Schultz, Zhenlin Ju, Bradley M. Broom, Alexander J. Lazar, A. Gordon Robertson, Mauro A. A. Castro, Jesper B. Andersen, Ioannis Tsamardinos, Vincenzo Lagani, Paulos Charonyktakis, Joshua M. Stuart, Christopher K. Wong, Verena Friedl, Toshinori Hinoue, Vladislav Uzunangelov, Peter W. Laird, Andrew D. Cherniack, Lindsay Westlake, Whijae Roh, Gad Getz, Stephanie H. Hoyt, Theo A Knijnenburg, Christina Yau, Jordan A. Lee, Lewis R. Roberts, Kyle Ellrott, Jasleen Grewal, Steven Jones, Chen Wang, Brian J Karlberg, Akinyemi I. Ojesina, Christopher C Benz, Kami E Chiotti, Katherine A. Hoadley, Ilya Shmulevich, Bahar Tercan, Galen F. Gao, Taek-Kyun Kim, Esther Drill, Ronglai Shen, Daniele Ramazzotti.

## Author Contributions

**WR** Conceptualization, software, formal analysis, validation, investigation, visualization, methodology, writing–original draft, writing–review and editing. **YG** Conceptualization, formal analysis, validation, investigation, visualization, methodology, writing–original draft, writing– review and editing. **MM** Writing–review and editing. **SA** Data curation. **JK** Software. **DH**, Data curation. **JFG** Writing–review and editing. **PWL** Resources. **ADC** Resources, data curation, formal analysis, writing–review and editing. **GG** Conceptualization, resources, data curation, supervision, funding acquisition, writing–review and editing. All authors read and approved the final manuscript.

## Acknowledgements

The Authors would like to thank The Cancer Genome Atlas Analysis Network (TCGA) and the Clinical Proteomic Tumor Analysis Consortium (CPTAC) for generating the data and making it available for use in our study. We thank Dr. Matthew Meyerson’s lab at the Broad institute for providing us all needed cell lines for this work.

