## Supplemental figures for "High-resolution lung adenocarcinoma expression subtypes identify tumors with dependencies on *MET, CDK4, CDK6*, and *PD-L1*"

**Supplementary Note 1. Concordance between our subtypes and intermediate mRNA expression clusters from Chen et al., 2016 study**

The concordance between our subtypes and intermediate mRNA expression clusters from Chen et al., 2016 study (4) was studied. We analyzed the intermediate clustering of mRNA expression data containing 6 clusters, which were significantly different from their final COCA subtypes (**Figure S1B, Table S2**). Indeed, when we compared these 6 mRNA-based clusters with our 5 mRNA–based subtypes, as well as the previously published PP, PI, and TRU expression-based subtypes, we observed a high concordance among them all (**Figure 1B Table S2**). The majority of LUAD samples (91%) were assigned to their clusters 4, 5, and 6, and these three clusters also mapped to the previously defined subtypes PP, PI, and TRU, respectively (**Figure S1C**). Comparing directly to our subtypes, 74.7% of our S4 tumors mapped to their cluster 4, while 83.7% of our S5 tumors mapped to their cluster 6. Moreover, their cluster 5 could be further partitioned into our S1, S2, and S3, which is consistent with cluster 5 mapping to the PI subtype (**Figure 1B, Table S2**).

In contrast to the concordance we observed when comparing our expression data to the interim expression data in Chen et al., 2016 study, there was a low concordance between our subtypes and their full COCA-based lung adenocarcinoma (AD) subtypes (**Figure S1A, Table S2)**. This low concordance is not surprising since the COCA subtypes were defined across all NSCLC (including both LSCC and LUAD) tumors and were obtained by a different procedure in which the tumors were first independently clustered based on different genomic features (DNA copy number, DNA methylation, mRNA expression, miRNA expression, and protein) and then the final clustering was based on their cluster assignments across the different features.

Supplementary Figure 1

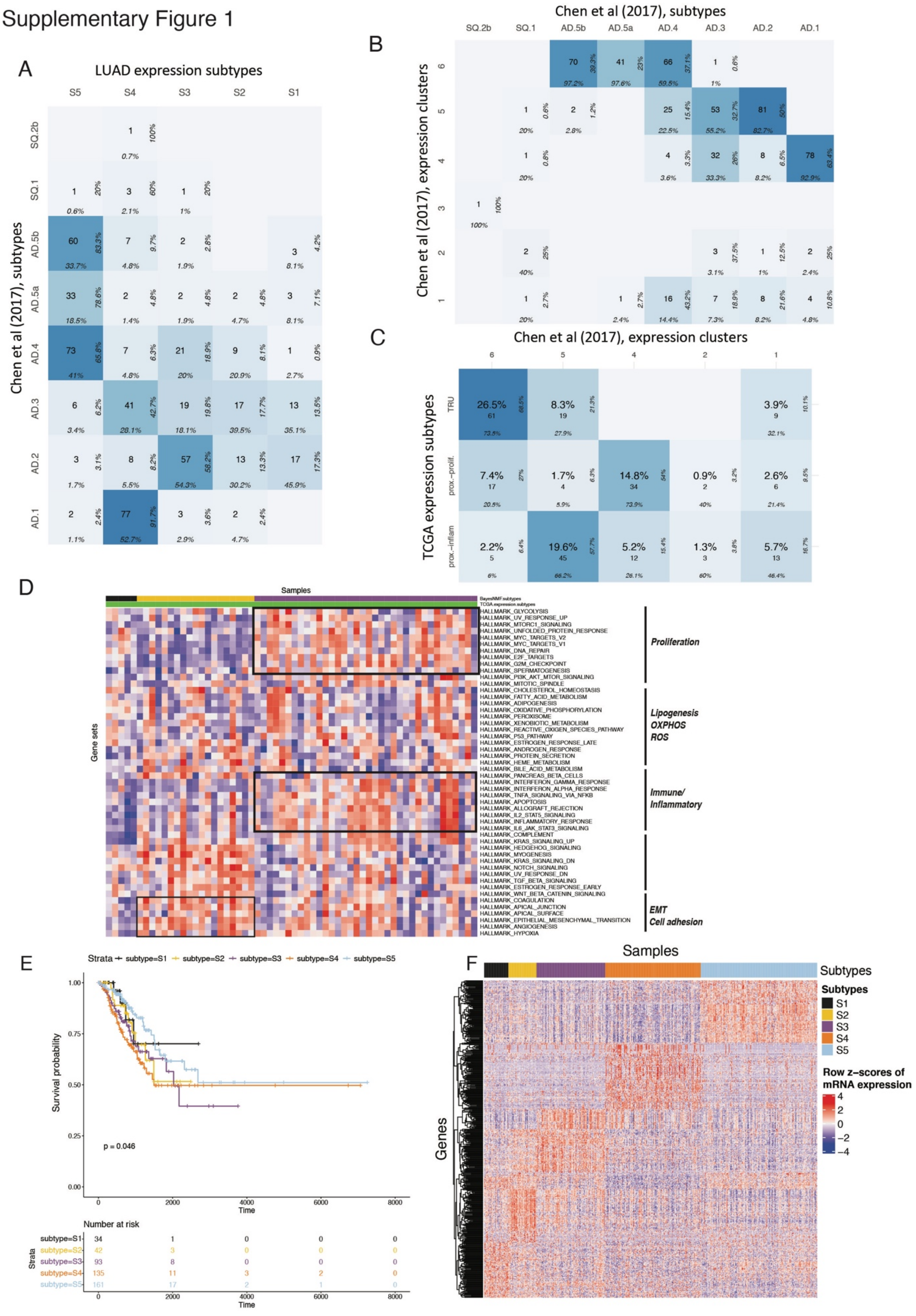

**Supplementary Figure 1. Identification and characterization of 5 newly identified LUAD expression subtypes**

(**A-C**) The confusion matrix shows concordance between two groups of subtypes of interest. The cell count in the middle shows the number of samples overlapping between two subtypes. The column-wise proportion is shown at the bottom of each cell and the row-wise proportion is shown on the right side of each cell. (**A**) LUAD expression subtypes comparison to the full Chen et al., 2017 (4) subtypes. (**B**) Full Chen et al., 2017 (4) subtypes comparison to their own expression subtypes. (**C**) Chen et al., 2017 (4) expression subtypes comparison to the TCGA expression subtypes.

(**D**) The heatmap shows overall pathway activation profiles (in row z-scores of GSVA enrichment scores for MSigDB hallmark gene sets) of tumors with PI subtype mapped to S1, S2 or S3.

(**E**) Kaplan-Meier curves for the disease-specific survival (DSS) among TCGA LUAD expression subtypes. The *P* value was calculated by the log rank test.

(**F**) The heatmap shows row z-scores of mRNA expression of subtype marker genes across 5 expression subtypes.

#### Supplementary Figure 2

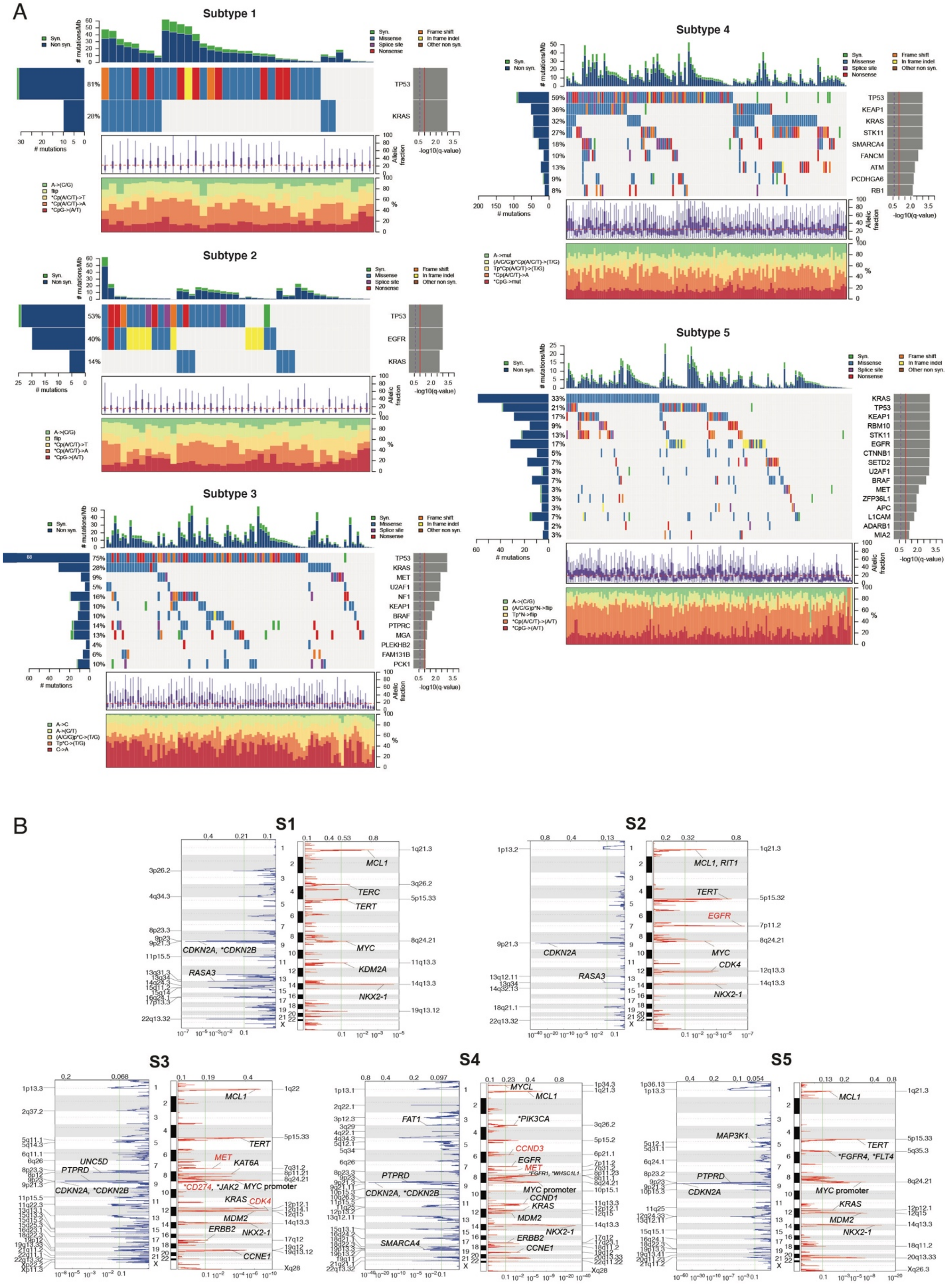

**Supplementary Figure 2. Co-mutation plots by subtypes (MutSig2CV)**

(**A**) MutSig2CV was run for each expression subtype for subtype-specific driver mutation discovery. The resulting co-mutation plots show the driver mutations for each subtype based on the Q value cutoff of 0.1.

(**B**) GISTIC2.0 was run for each expression subtype to define subtype-specific recurrent SCNA. The resulting GISTIC plots show the recurrent SCNAs for each subtype based on the Q value cutoff of 0.1.

Supplementary Figure 3

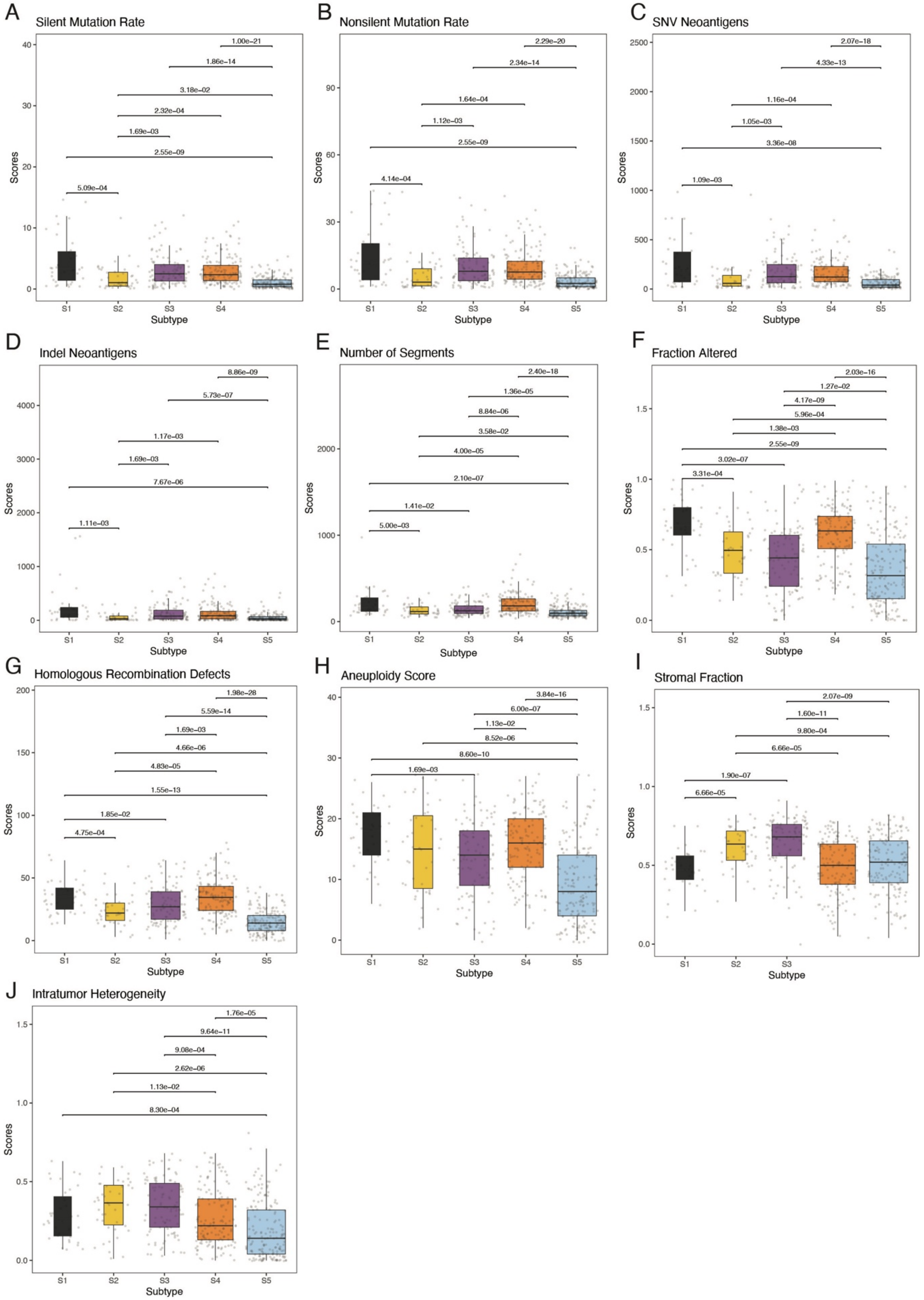

**Supplementary Figure 3. Additional subtype-specific genomic feature data obtained from Thorsson et al., 2018**

The boxplots (**A-J**) show the scores of genomic features obtained from Thorsson et al., 2018 (24) among LUAD expression subtypes.

Supplementary Figure 4

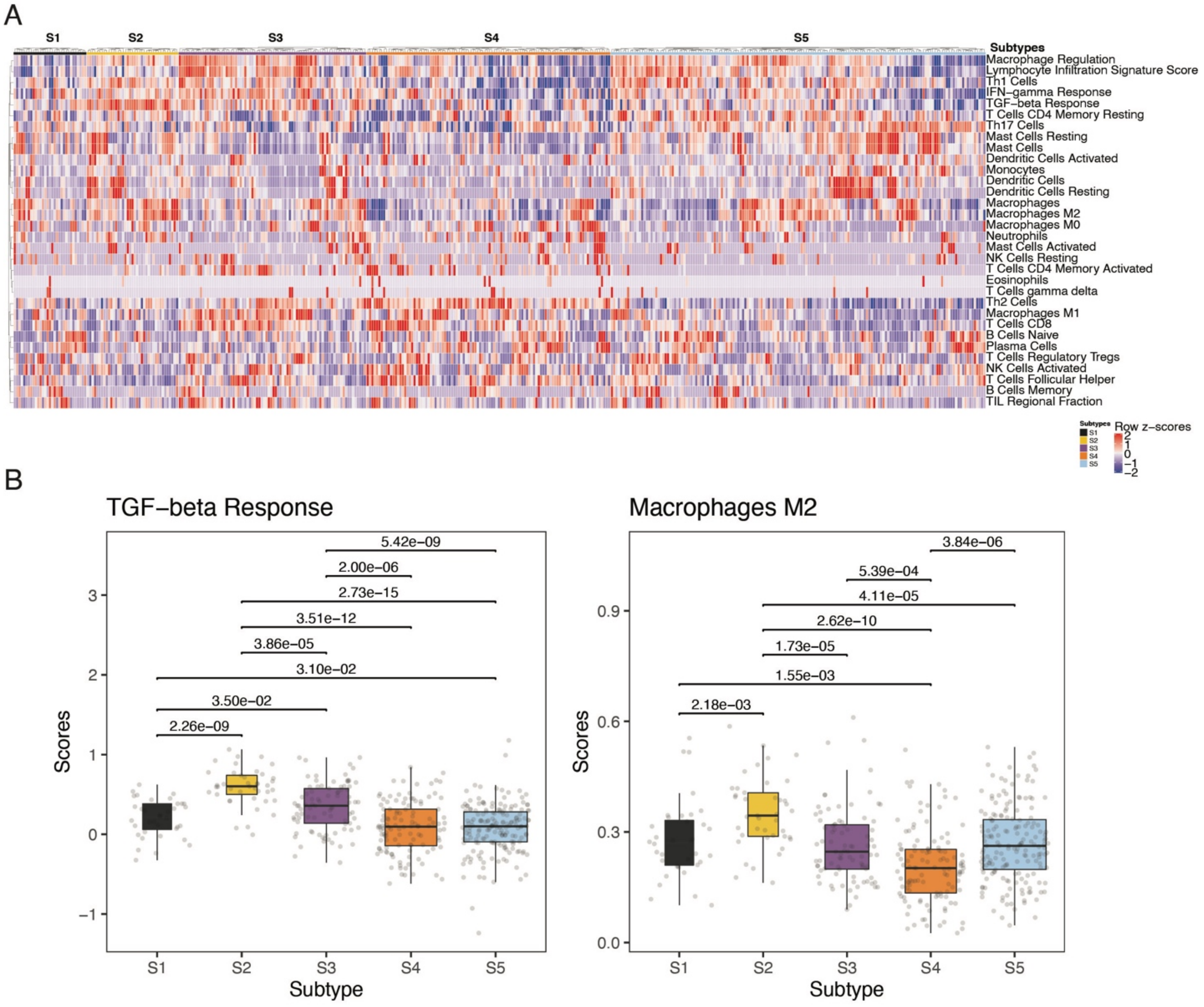


**Supplementary Figure 4. Additional subtype-specific immune cell subset fraction data obtained from Thorsson et al., 2018**

(**A**) The heatmap shows immune cell subset fraction (in row z-scores of immune cell subset fraction values obtained from Thorsson et al., 2018 (24)) of TCGA LUAD tumors across expression subtypes.

(**B**) The boxplots show selected immune cell subset fraction (obtained from Thorsson et al., 2018 (24)) among LUAD expression subtypes.

### Supplementary Figure 5

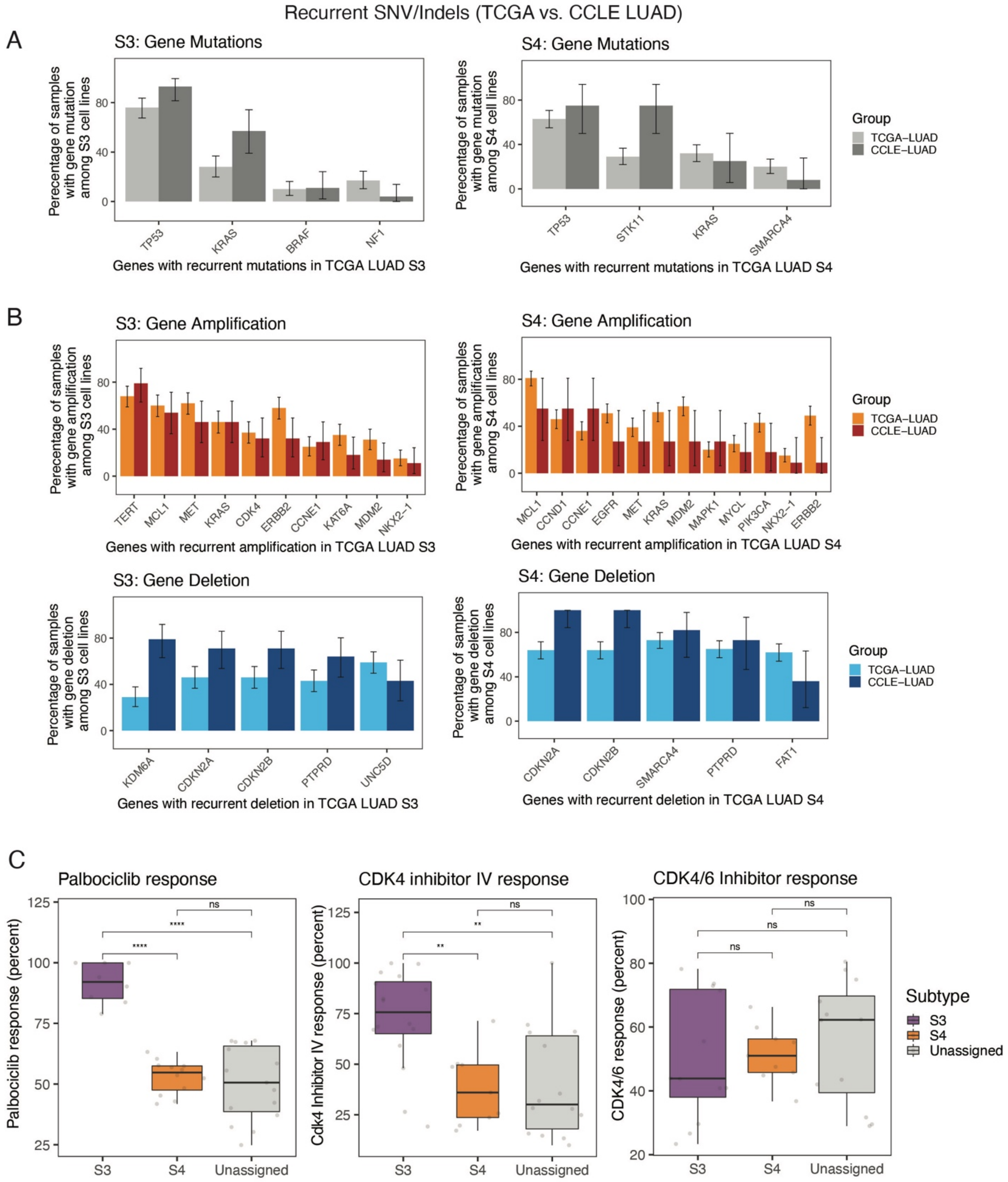

**Supplementary Figure 5. Concordance of frequencies in recurrent genomic alterations for S3 and S4 between TCGA and CCLE LUAD expression subtypes**

(**A**) The histogram shows the percentage of TCGA/CCLE LUAD samples with recurrent SNVs/Indels observed in the TCGA LUAD cohort. The Bayesian credible interval was used for a 95% credible interval.

(**B**) The histogram shows the percentage of TCGA/CCLE LUAD samples with recurrent copy number alterations observed in the TCGA LUAD cohort. The Bayesian credible interval was used for a 95% credible interval.

(**C**) The boxplots show response to CDK4/6 inhibitors measured by the delta change in confluency between treated and untreated (DMSO only) cell lines within each subtype. Left panel - response to Palbociclib (CDK4 specific concentration - 11nM) , middle panel - response to CDK4/6 Inhibitor IV (CDK4 specific concentration - 1.5 μM) and right panel - response to Palbociclib (CDK4/6 concentration - 16 nM).

Supplementary Figure 6

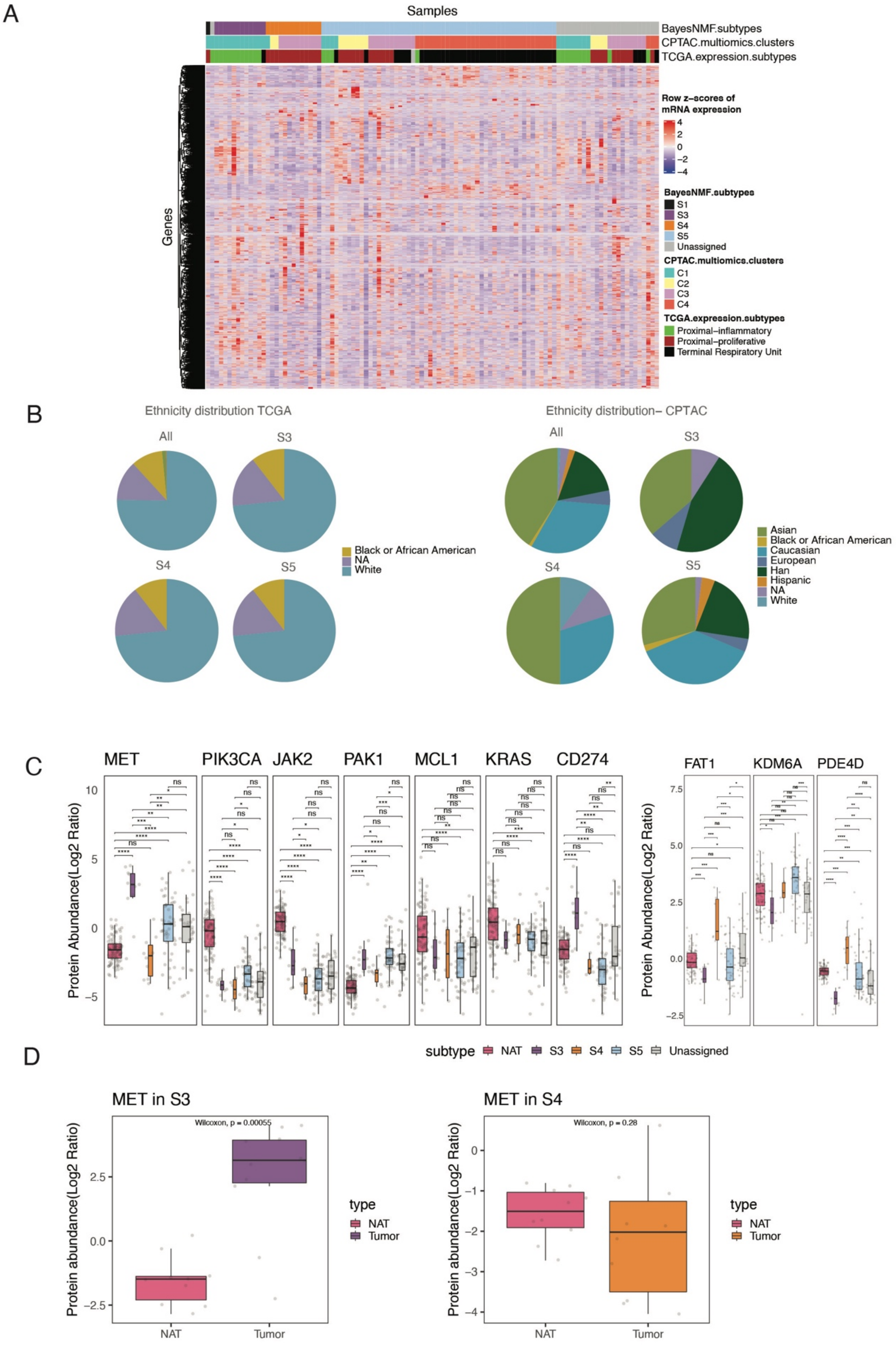

**Supplementary Figure 6. CPTAC LUAD expression subtypes and protein expression of genes with recurrent SCNAs in TCGA LUAD**

(**A**) The heatmap shows row z-scores of mRNA expression of 5,000 most variable genes across CPTAC LUAD samples. The upper column annotations show the newly identified TCGA expression subtypes (upper), CPTAC multi-omics clusters (middle), and the original TCGA LUAD expression subtypes (lower) of CPTAC LUAD samples.

(**B**) Ethnicity distribution across all, S3, S4, or S5 tumors in TCGA LUAD cohort versus CPTAC LUAD cohort

(**C**) The boxplots show protein abundance of genes with recurrent SCNAs in tumors versus normal adjacent tissues among CPTAC LUAD expression subtypes.

(**D**) The boxplots show *MET* protein expression in NAT versus tumors in S3 and S4.

Supplementary Figure 7

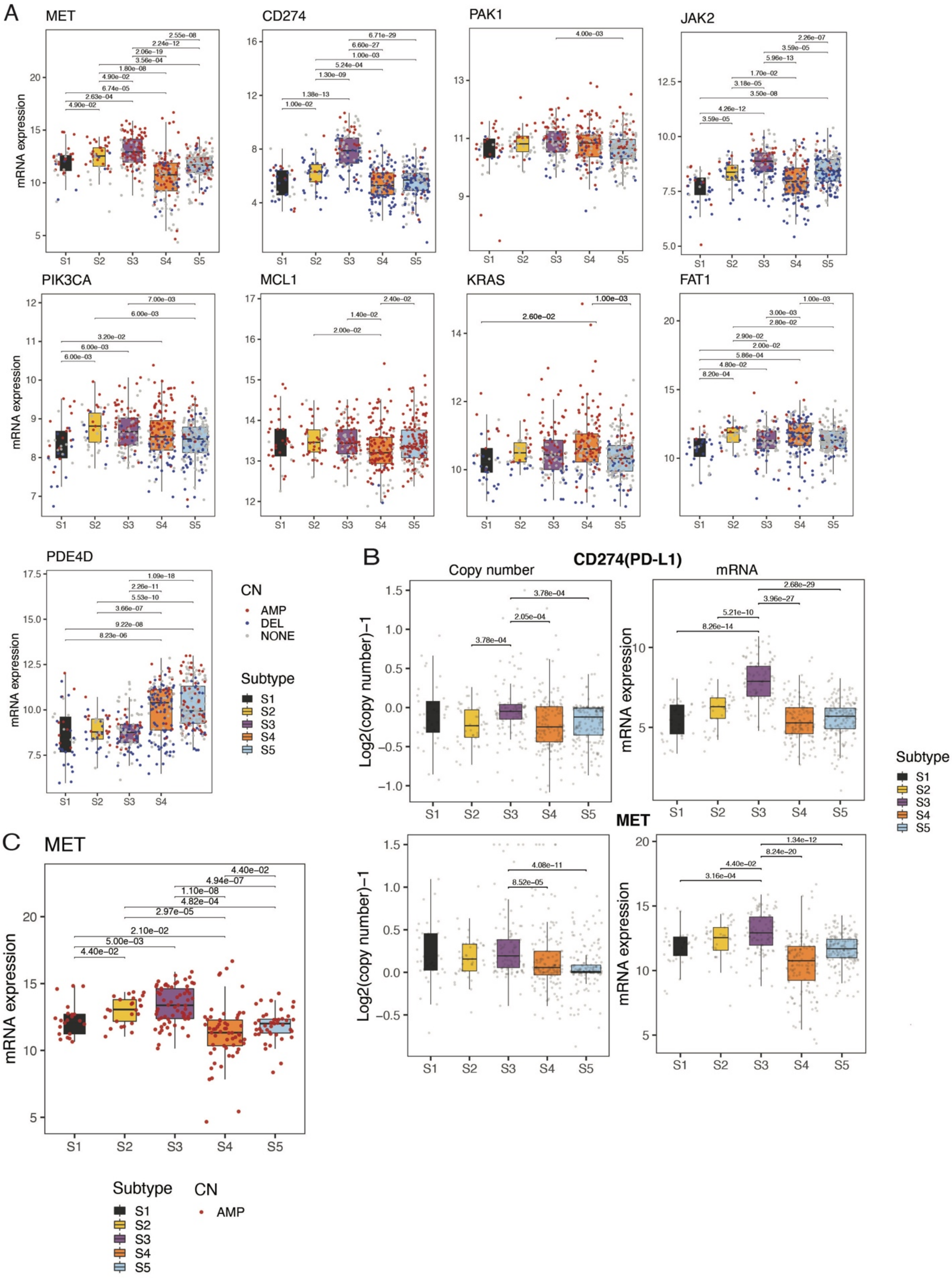

**Supplementary Figure 7. Expression analysis of genes with subtype-specific recurrent SCNAs**

(**A**) The boxplots show mRNA expression of genes with recurrent SCNAs across LUAD expression subtypes. Copy number states of genes are shown with different colors (red: amplification, blue: deletion, gray: no SCNAs).

(**B**) The boxplots show copy number and mRNA expression of *CD274* (PD-L1) gene - upper panel and *MET* gene- lower panel across LUAD expression subtypes in TCGA data.

(**C**) The boxplots show MET mRNA expression comparison across LUAD expression subtypes when restricting the analysis to only MET-amplified tumors.

Supplementary Figure 8

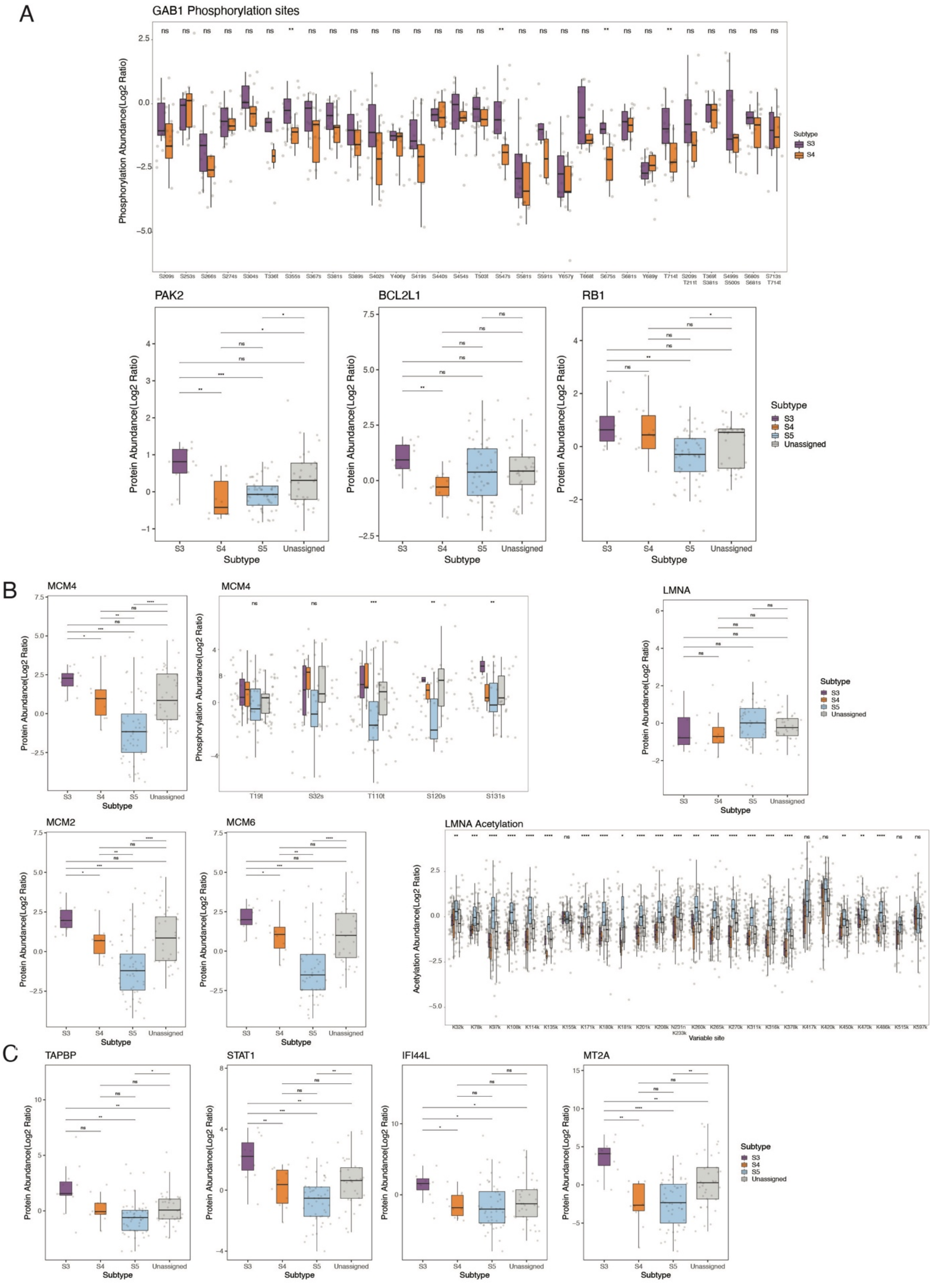

**Supplementary Figure 8. Proteomic evidence of cell proliferation and immune pathway activation**

(**A**) The boxplots show phosphorylation level and protein abundance of genes supporting *MET* pathway activation in S3 versus S4.

(**B**) The boxplots show phosphorylation level and protein abundance of genes involved in cell proliferation among CPTAC LUAD expression subtypes.

(**C**) The boxplots show phosphorylation level and protein abundance of genes involved in antigen presentation and interferon signaling among CPTAC LUAD expression subtypes.

**Supplementary Tables (please see attached Excel files)**

**Supplementary Table S1. TCGA LUAD sample-subtype association scores**

The table shows the normalized association scores (in the normalized H matrix) of each TCGA LUAD sample to each of the five LUAD expression subtypes. Samples were assigned to one of the five identified LUAD expression subtypes if the normalized association with the subtype was larger than 0.6. Otherwise, samples were annotated as 'Unassigned'.

**Supplementary Table S2. Subtype membership of TCGA LUAD samples**

The table shows the subtypes membership of each TCGA LUAD sample to multiple subtype definitions (columns: 'Subtype' - our subtypes, 'COCA' - COCA subtypes, 'COCA_expression_cluster' - expression clusters used in the COCA clusters, 'TCGA_subtype' - expression subtypes used in the TCGA LUAD paper).

**Supplementary Table S3. Differentially-regulated pathways in LUAD expression subtypes**

The table sheet 1 ('All_LUAD_up_down_pathways') shows the differentially-regulated pathways (MSigDB hallmark gene sets) for each LUAD expression subtype. The table sheet 2 ('PI_split_up_and_down_pathways') shows the differentially-regulated pathways for 60 tumors that originally were assigned to the PI subtype and were further partitioned in our S1, S2 and S3 subgroups. Significant differentially-regulated pathways were defined as the pathways with Q value < 0.05 and mean difference in GSVA enrichment scores between subtype of interest and others > 0.2 or < -0.2. *P* values were calculated by the Wilcoxon rank sum test. *Q* values are FDR-adjusted *P* values.

**Supplementary Table S4. CCLE LUAD sample-subtype association scores**

The table shows the normalized association scores (in the normalized H matrix) of each CCLE LUAD cell line to each of the five LUAD expression subtypes. The cell lines were assigned to one of the five identified LUAD expression subtypes if the normalized association with the subtype was larger than 0.6. Otherwise, samples were annotated as 'Unassigned'.

**Supplementary Table S5. Subtype marker gene list**

The table shows the subtype marker genes for each of the five LUAD expression subtypes.

**Supplementary Table S6. S3/S4-specific cancer vulnerabilities**

The tables show the significant S3-associated cell line-specific cancer vulnerabilities (in sheet 1: "S3_vs_Others") and the S4-associated cell line-specific cancer vulnerabilities (in sheet 2: "S4_vs_Others"). The table sheet 3 lists the Achilles common essential genes that were removed from analysis.

**Supplementary Table S7. CPTAC LUAD sample-subtype association scores**

The table shows the normalized association scores (in the normalized H matrix) of each CPTAC LUAD sample to each of the five LUAD expression subtypes. Samples were assigned to one of the five identified LUAD expression subtypes if the normalized association with the subtype was larger than 0.6. Otherwise, samples were annotated as 'Unassigned'.

**Supplementary Table S8. Biomarkers for S3 and S4 tumors**

The table shows the biomarker discovery analysis by lasso logistic regression models. '[S3/S4]_gene_expression' tab shows the results from S3/S4 prediction models based on TCGA LUAD gene expression data. '[S3/S4]_gene_expression_lambda' tab shows the accuracy of the S3/S4 prediction models based on TCGA LUAD gene expression data with varying lambda values. '[S3/S4]_RPPA' tab shows the results from S3/S4 prediction models based on TCGA LUAD RPPA data. '[S3/S4]_RPPA_lambda' tab shows the accuracy of the S3/S4 prediction models based on TCGA LUAD RPPA data with varying lambda values.
